# Metabolic diversity in commensal protists regulates intestinal immunity and trans-kingdom competition

**DOI:** 10.1101/2022.08.26.505490

**Authors:** Elias R. Gerrick, Soumaya Zlitni, Patrick T. West, Matthew M. Carter, Claire M. Mechler, Matthew R. Olm, Elisa B. Caffrey, Jessica A. Li, Steven K. Higginbottom, Christopher J. Severyn, Frauke Kracke, Alfred M. Spormann, Justin L. Sonnenburg, Ami S. Bhatt, Michael R. Howitt

**Author notes:** These authors contributed equally (Soumaya Zlitni, Patrick T. West, Matthew M. Carter).

## Abstract

The microbiota influences intestinal health and physiology, yet the contributions of commensal protists to the gut environment have been largely overlooked. Here, we identified several new rodent- and human-associated parabasalid protists. Genomic and metabolomic analyses of murine parabasalids from the genus *Tritrichomonas* revealed species-level differences in the excretion of the metabolite succinate. This metabolic dissimilarity results in distinct small intestinal immune responses during protist colonization. Metabolic differences between *Tritrichomonas* species also determine their ecological niche within the microbiota. By manipulating dietary fibers and developing *in vitro* protist culture, we show that different parabasalid species preferentially rely on dietary polysaccharides or mucus glycans. These polysaccharide preferences create trans-kingdom competition with specific commensal bacteria, which affects intestinal immunity in a diet-dependent manner. Our findings reveal unappreciated diversity in commensal parabasalids, elucidate differences in commensal protist metabolism, and suggest how dietary interventions could regulate their impact on gut health.

## Introduction

The gut microbiota is a complex and diverse microbial ecosystem that broadly impacts the development and function of the intestinal immune system^1–5^. This relationship has been studied predominantly with microbial communities, where conserved microbial ligands and metabolites have pleiotropic effects on the host^6–10^. However, some microbial species dominantly influence the immune system independent of other microbiota members^11–13^. For example, segmented filamentous bacteria induce T helper 17 (Th17) cells in the murine small intestine, and *Bacteroides fragilis* stimulates regulatory T cells in the colon^12,13^. While bacteria are the most well-characterized immunologically dominant symbionts, the gut microbiota also contains viruses, fungi, archaea, and protists that interact with the host and alter immune function.

Protists are often overlooked as members of the microbiota, in part due to the historical view of these microbes as universally pathogenic. However, commensal protists, including those in the *Parabasalia* phylum, are present in healthy mammalian guts and can dominantly shape intestinal immunity in mice^14–18^. Parabasalid protists from the genus *Tritrichomonas* are widespread in laboratory mice and dramatically alter the gut mucosa^15,19–21^. In the small intestine, *Tritrichomonas* species drive type 2 immunity through production of the metabolite succinate, which lengthens the gut and promotes tuft and goblet cell hyperplasia in the epithelium^20,21^. In the colon, *Tritrichomonas musculis* (*Tmu*) activates the inflammasome to promote Th1 and Th17 immunity^15,22^. The effects of these tritrichomonads on epithelial and immune functions significantly alter the severity of some enteric infections and several inflammatory diseases in mouse models, and do not induce apparent pathology^15,20–22^. In humans, only two species of commensal parabasalids have been identified, although their effects on intestinal immunity are still unclear. Thus, despite the growing appreciation of how commensal parabasalids impact mucosal tissues, very little is understood about their diversity, metabolism, or microbial ecology.

Fundamental aspects of commensal parabasalid protist biology have been obscured by a lack of genomic information and their resistance to *in vitro* culture. Consequently, the biology of commensal *Tritrichomonas* species is largely inferred from studies on the urogenital pathogenic parabasalids *Trichomonas vaginalis* and *Tritrichomonas foetus.* However, these pathogens infect different hosts and anatomical niches that may not reflect the specific adaptations of commensal parabasalids in the gut. Additionally, the ecological role of commensal parabasalids within the mammalian gut microbiota is unexplored. Parabasalid protists are generally fermentative^21,23–25^, but the specific carbohydrate sources used by commensal species and how their harvesting of these carbohydrates affects the intestinal environment is unknown. Thus, parabasalid metabolism may contain unidentified diversity that shapes commensal protists’ interactions with both the host and microbiota.

In this work, we identified parabasalids in humans from both industrialized and non-industrialized populations, revealing large amounts of unappreciated diversity in human-associated parabasalids. In addition, we identified a novel murine *Tritrichomonas* species called *Tritrichomonas casperi.* Using genomic, transcriptomic, and metabolomic approaches, we determined that fermentative output from *T. casperi* differs from previously characterized murine tritrichomonads like *Tmu,* which results in divergent immune responses in the small intestine. Further, we used this metabolic information to create a framework for predicting the production of immunomodulatory metabolites by parabasalids, including human-associated species, based only on phylogeny. Next, we demonstrated that these different species’ preferences for fermenting plant polysaccharides or mucus glycans affect trans-kingdom interactions with bacteria. Finally, through highly controlled studies on carbohydrate utilization, we revealed that the composition of dietary fiber acts as a lever to control protist-induced type 2 immunity. Our work provides mechanistic insights into how commensal parabasalids, through their metabolic differences, act as architects of the intestinal ecosystem and establishes a foundation for further mechanistic studies of how these protists influence host immunity and microbiome ecology.

## RESULTS

### Discovery of a new commensal *Tritrichomonas* species in mice

To discover uncharacterized parabasalid protists within laboratory mice, we screened DNA extracted from stool using pan-parabasalid PCR primers. These primers bind to highly conserved sequences of parabasalid rRNA loci and generate internally transcribed spacer (ITS) amplicons sufficient to distinguish parabasalid species using Sanger Sequencing. As expected, we identified *Tmu* (previously referred to as *T. muris*^19^) in a subset of our mice, but stool from another group of mice produced mixed signals that indicated the presence of multiple closely related parabasalid species. Examination of the cecal contents of these mice by light microscopy revealed protists with two distinct sizes: a large variant consistent in size with *Tmu* and another variant approximately half the length.

To determine if these two sizes constitute different morphologies of *Tmu* or two separate species, we purified both sizes of protist by fluorescence-activated cell sorting (FACS) and colonized protist-negative specific pathogen free (SPF) mice with either the large or small variant (Figure 1A). Mice maintained colonization with homogenously small or large protists, as verified by scanning electron microscopy (SEM), suggesting these were separate species (Figure 1B and S1A). Ultrastructural examination of both protists revealed an undulating membrane and recurrent flagellum typical of the *Parabasalia* phylum (Figure 1B)^26^. Additionally, the new protist, like *Tmu,* had three anterior flagella, suggesting it belongs to the genus *Tritrichomonas*. To confirm this phylogenetic relationship, we compared ITS sequences from both protists, which revealed the smaller species is an uncharacterized murine isolate with accession number MF375342 that shares 83.4% identity with *Tmu* (Figure S1B). Thus, the smaller protist is a new species, hereafter referred to as *Tritrichomonas casperi* (*Tc*). Healthy mice maintained high levels of *Tc* colonization for at least 3 months, indicating that *Tc* is a stable member of the murine microbiota (Figure S1C).

**Figure 1.**
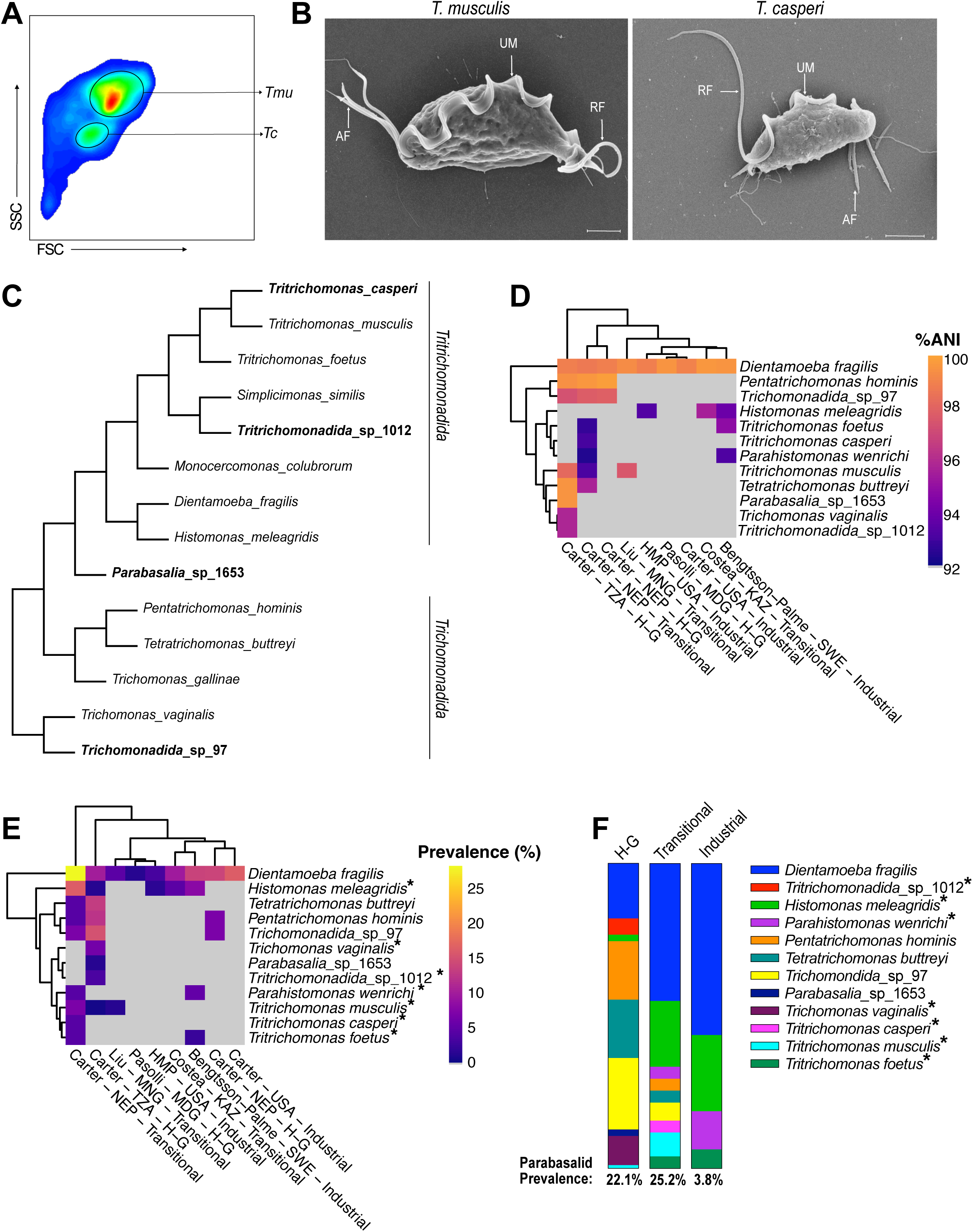
Identification of novel parabasalids in mice and humans. (A) FACS purification of the large *T. musculis* (*Tmu*) and small *T. casperi* (*Tc*) protists isolated from a mouse cecum. FSC, forward scatter. SSC, side scatter. (B) SEM images of *Tmu* (left) and *Tc* (right). UM: Undulating Membrane, RF: Recurrent Flagellum, AF: Anterior Flagella. Scale bars are 2µm. (C) Cladogram of parabasalid protists based on ITS sequences, including novel mouse and human-associated parabasalids. Protists in bold indicate novel parabasalids discovered in this study. (D-F) Identification of human-associated parabasalids using mapping-based analysis of metagenomic data. (D) Percentage average nucleotide identity (ANI) of parabasalid metagenomic reads mapping to the corresponding reference protist sequence. Imperfect matches indicate that identified parabasalids are relatives of reference protists and not the same species. First author of the cohort study, country of origin, and population type (“Industrial”, “Transitional”, Hunter-Gatherer (“H-G”)) is indicated. Columns and rows are hierarchically clustered based on Euclidean distance. Cells corresponding to protists not identified in a population are colored grey. (E) Percentage prevalence of subjects in human cohorts with metagenomic reads mapping to reference parabasalids. (F) Prevalence of parabasalids identified in each population type. Asterisks denote protists with imperfect matches to the reference sequence, which are therefore relatives of the indicated species (E, F). (Abbreviations: TZA=Tanzania, NEP=Nepal, MNG=Mongolia, USA=United States of America, MDG=Madagascar, KAZ=Kazakhstan, SWE=Sweden, HMP=Human Microbiome Project).

### Characterization of parabasalid diversity in humans

Rodent *Tritrichomonas* species are not known to colonize humans, but two commensal parabasalid species, *Pentatrichomonas hominis* and *Dientamoeba fragilis*, can be found in the human gut. However, most studies have used targeted approaches that are unlikely to detect diverse species and primarily investigate samples from industrialized populations which harbor less diverse microbiomes^18,27,28^. Therefore, to identify human-associated commensal parabasalids we created a reference set of 29 parabasalid ITS sequences (Table S1) to query previously published metagenomes from 1,800 human fecal samples representing 29 different populations^28^. These metagenomes encompass a wide array of human lifestyles, including industrial and hunter-gatherer populations, as well as “transitional” populations which have neither hunter-gatherer nor industrialized lifestyles. Using our parabasalid ITS database as reference sequences, we assembled ITS sequences from five parabasalids in these human samples, as well as a novel *Simplicimonas* species from cow feces which we refer to as *Tritrichomonadida_*sp_1012. While the human-associated sequences included *D. fragilis* and *P. hominis*, they also contained *Tetratrichomonas buttreyi*, which has only previously been identified in cattle. We also identified two previously undescribed candidate parabasalid species in the microbiomes of non-industrialized populations, which we refer to as *Trichomonadida*_sp_97 and *Parabasalia*_sp_1653 (Figure 1C and Table S1). Phylogenetic analysis suggested that *Trichomonadida_*sp_97 belongs to the *Trichomonadida* order and *Parabasalia*_sp_1653 potentially represents a novel *Parabasalia* order (Figure 1C).

The method used to identify these six parabasalids in the metagenomic assemblies suffers from low sensitivity inherent to assembly-based approaches. This assembly-based sensitivity issue is further compounded by low sequencing depth in a subset of the metagenomic datasets which may have limited the number of identified protists. We therefore analyzed these datasets using a mapping-based approach, which improves sensitivity of detection but does not enable accurate classification beyond identifying the closest match in our reference database (Figure S1D and 1E). This mapping-based methodology revealed a strong effect of sequencing depth on parabasalid identification; we therefore implemented a 5Gbp minimum sequencing depth threshold for further analysis (Figure S1F). In this curated database of 557 metagenomes, there was an inverse relationship between industrialization and parabasalid colonization frequency, with parabasalids identified in more than 22% of samples from non-industrialized populations but only 3.8% of industrialized samples (Figure 1D, 1E, 1F). *D. fragilis* was the most prevalent parabasalid in industrialized and transitional populations, but not in the hunter-gatherer populations. The microbiomes of non-industrialized individuals contained many other parabasalid species including *T. buttreyi*, *Trichomonadida*_sp_97, and *Parabasalia_*sp_1653. Similarly, parabasalids closely related to *T. vaginalis*, *Tritrichomonadida_*sp_1012, and the murine tritrichomonads *Tmu* and *Tc,* were also present in some non-industrialized populations (Figure 1D, 1E, and 1F). Together, these results demonstrate that a large amount of unappreciated diversity exists among human-associated commensal parabasalids and are consistent with previous studies suggesting that industrialization is associated with decreased diversity of microeukaryotes in the microbiome^17,18^.

### *Tmu* and *Tc* stimulate divergent immune responses in the small intestine

*D. fragilis* belongs to the same phylogenetic order as the murine tritrichomonads *Tmu* and *Tc*, and we identified even closer relatives of the murine protists in non-industrialized human populations. Thus, our results reveal new parabasalid diversity within human microbiomes and reinforce the utility of murine tritrichomonads for understanding the functional impact of commensal parabasalids on the intestinal environment. All previously characterized murine commensal tritrichomonads, including *Tmu*, induce tuft cell hyperplasia and type 2 immunity in the distal small intestine (SI) through excretion of the metabolite succinate^19–22^. However, tuft cell hyperplasia, which is used as a readout for SI type 2 immunity^19–21^, was absent in mice colonized with *Tc* (Figure 2A, 2B, 2C, and S2A). Although *Tc* stimulated a subtle increase in group 2 innate lymphoid cells (ILC2s), which are more sensitive to weak type 2 stimuli, this increase was markedly lower than the ILC2 expansion associated with *Tmu* (Figure 2D, 2E, and S2B). Moreover, *Tc* colonization reduced the proportion of small intestinal GATA3^+^ CD4^+^ T helper 2 (Th2) cells associated with adaptive type 2 immunity, which contrasted with the expansion of Th2 cells in *Tmu* colonized mice (Figure 2F and S2C). To determine whether *Tc* failed to drive type 2 immunity because of colonization deficiencies, we compared *Tc* and *Tmu* abundance by quantitative PCR. *Tc* was at least as abundant as *Tmu* along the length of the intestine, including the distal SI (Figure S2D). These results suggest that *Tc* is unique among known murine commensal tritrichomonads because it does not induce type 2 immunity in the distal SI.

**Figure 2.**
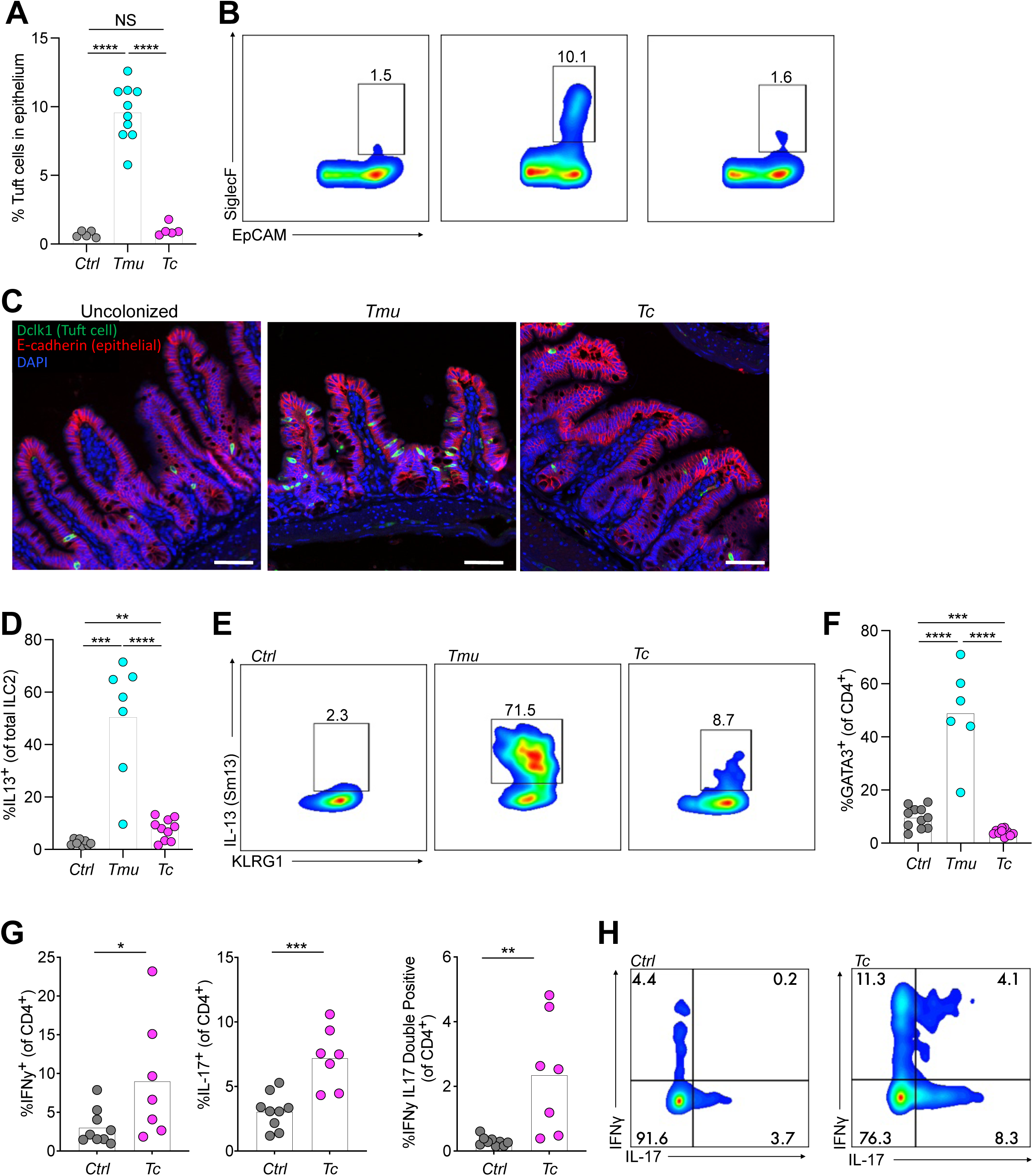
T Cell responses to *Tmu* and *Tc* differ in the distal SI. (A) Tuft cell frequency measured by flow cytometry in the distal SI epithelium of uncolonized (*Ctrl*) mice or those colonized with *Tmu* or *Tc* for two weeks. (B) Representative flow cytometry plots of tuft cells. SiglecF marks tuft cells, EpCAM marks epithelial cells. (C) Representative immunofluorescence microscopy images of tuft cell hyperplasia in the distal SI of mice with or without each protist for two weeks. Scale bars are 50µm. (D) Frequency of ILC2s in the lamina propria of the distal SI in *Ctrl* mice or mice colonized with *Tmu* or *Tc* for two weeks. IL13 was measured using the IL13 reporter Smart13 mice (E) Representative flow cytometry plots of ILC2s. (F) Frequency of GATA3^+^ CD4^+^ T cells (Th2 cells) in the lamina propria of the distal SI after three weeks of colonization. (G) Frequency of IFNγ and IL-17 positive CD4^+^ T cells in the distal SI lamina propria after three weeks of *Tc* colonization, or without protist colonization (*Ctrl*). Cytokines were stimulated *ex vivo* and cells were stained for intracellular cytokines. (H) Representative flow cytometry plots of Th1 and Th17 cells in the distal SI with and without *Tc*. Graphs (A, D, F, G) depict mean. Each symbol represents an individual mouse from two to three pooled experiments (A, D, F, and G). *p<0.05, **p<0.01, ***p<0.001, ****p<0.0001, NS, not significant by Student’s t test.

Previous work demonstrated that *Tmu* increases CD4^+^ T helper 1 (Th1) and T helper 17 (Th17) cells in the cecum and colon^15,22^. Therefore, we examined whether *Tc* stimulates these same immune responses in the distal SI. IFNγ producing Th1 cells and IL-17 producing Th17 cells, as well as IFNγ and IL17 double positive CD4^+^ T cells, significantly increased in the distal SI of *Tc* colonized mice compared to protist-free controls (Figure 2G, 2H, and S2C). Because *Tmu* stimulates Th2 cells in the distal SI, we wondered whether co-colonization would alter the mucosal response to these protists. To address this question, we co-colonized mice with *Tmu* and *Tc* or singly with each protist. Tuft cell hyperplasia was identical in co-colonized and *Tmu* singly colonized mice, and *Tc* and *Tmu* abundance was unaffected by co-colonization (Figure S2E and S2F). This finding suggests that *Tmu* dominates the mucosal immune response when both species are present simultaneously in the distal SI.

### Th1 and Th17 induction is a common tritrichomonad immune response

Although *Tmu* induces Th1 and Th17 cells in the colon and type 2 immunity in the distal SI, no study has examined the immune response in both intestinal tissues using the same protist isolate. This leaves some ambiguity regarding whether different strains or species contribute to these heterogeneous *Tmu*-induced immune responses. We therefore characterized CD4^+^ T cell populations induced by *Tmu* and *Tc* in the colon. Both *Tmu* and *Tc* significantly increased Th1, Th17, and IFNγ-IL17 double positive T cells in SPF and gnotobiotic mice (Figure 3A, 3B, and S3A). Other T cell responses appeared unaffected by protist colonization; for example, regulatory T cells did not change with colonization by either species (Figure S3B).

**Figure 3.**
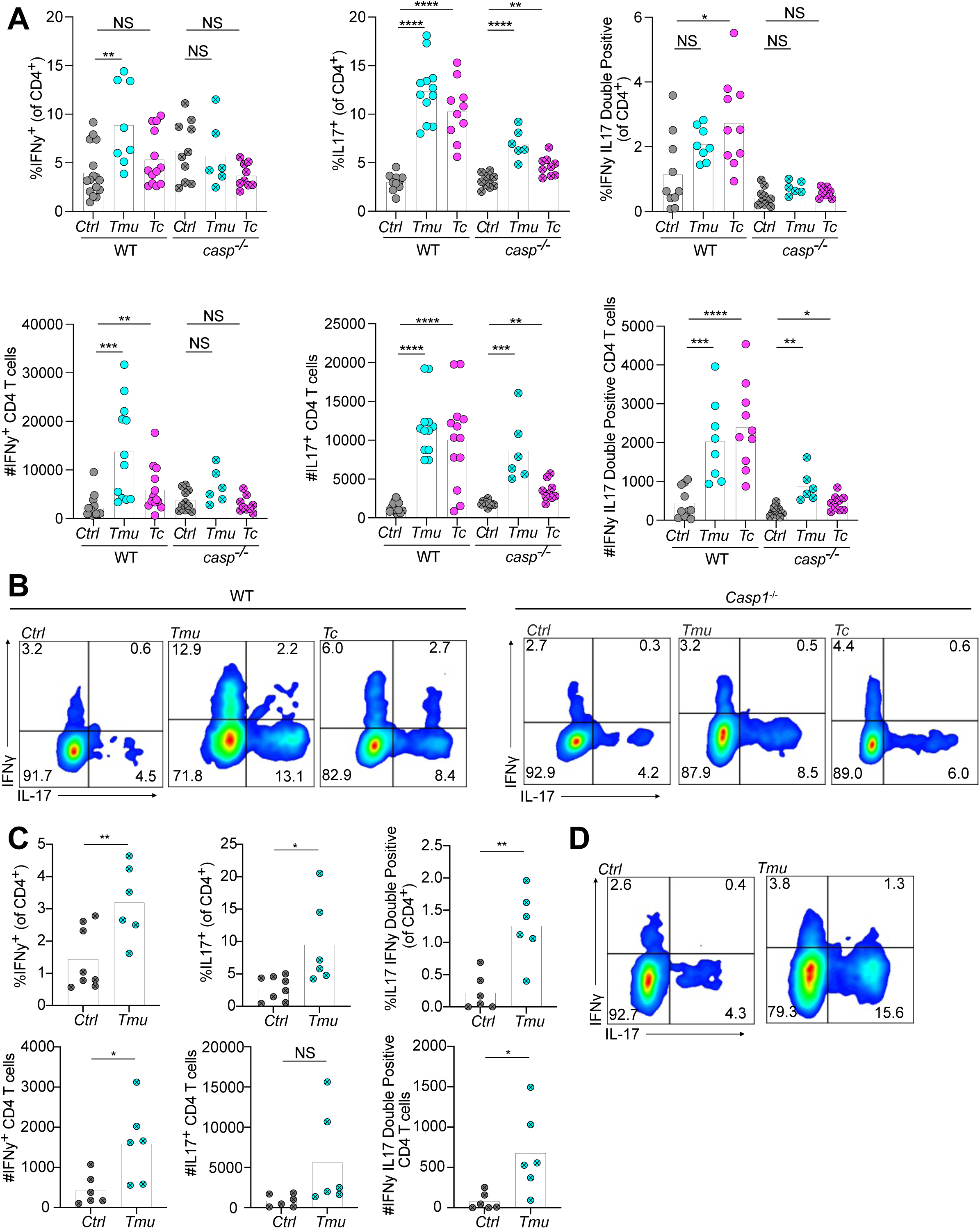
Th1 and Th17 induction is the shared tritrichomonad-induced immune response. (A) Frequency (top) and absolute abundance (bottom) of IFNγ and IL-17 positive CD4^+^ T cells in the colonic lamina propria of WT and *Caspase1^-/-^* mice after three weeks of protist colonization, or without protist colonization (*Ctrl*). (B) Representative flow cytometry plots of the frequency of IFNγ and IL17 positivity in colonic lamina propria CD4^+^ T cells. (C) Frequency and absolute abundance of IFNγ and IL-17 positive CD4^+^ T cells in the distal SI lamina propria of *Trpm5*^-/-^ mice after three weeks of *Tmu* colonization, or without protist colonization (*Ctrl*). (D) Representative flow cytometry plots of the frequency of IFNγ and IL17 positivity in distal SI lamina propria CD4^+^ T cells from *Trpm5^-/-^* mice with or without *Tmu*. Cytokines were stimulated *ex vivo* and cells were stained for intracellular cytokines. Graphs depict mean. Each symbol represents an individual mouse from two (C) or three (A) pooled experiments. *p<0.05, **p<0.01, ***p<0.001, ****p<0.0001, NS, not significant by Student’s t test.

Microbial adhesion has been proposed as a nonspecific Th17-inducing signal^29^, and previous work identified examples of *Tmu* adhering to colonic epithelial cells^15^. To address the prevalence of epithelial adhesion by *Tmu* and *Tc,* we developed a polyclonal antibody against *Tmu*. This antibody recognizes *Tmu,* and to a lesser extent *Tc,* and stains both trophozoites and pseudocysts of *Tmu* (Figure S3C and S3D). We gently flushed the colons of *Tmu* colonized mice with PBS to remove unadhered protists and then searched for adherent protists by fluorescence microscopy. Adherent protists were extremely rare (0-3 per colonic tissue section) and thus are unlikely to drive immune responses across the entire tissue (Figure S3E, S3F, and S3G).

*Tmu* requires the inflammasome to fully induce colonic Th1 and Th17 cells^15^, so we hypothesized that *Tc* also stimulates Th1 and Th17 immunity by activating the inflammasome. Mice deficient in the inflammasome component Caspase 1 had partially suppressed colonic immune induction by *Tmu* (Figure 3A and 3B)^15^. Caspase 1 was similarly important for the immune response in *Tc* colonized mice, suggesting that both protists stimulate Th1 and Th17 immunity in the colon through a shared effector.

Although *Tmu* and *Tc* induce the same Th1 and Th17 immune responses in the colon, only *Tc* appears to drive this response in the distal SI. Induction of Th1 and Th17 immunity in the distal SI by *Tc* demonstrates that the host pathway is present. Therefore, we reasoned that the lack of Th1 and Th17 induction in the distal SI by *Tmu* may be due to succinate driven type 2 immunity masking this response. The poor cellular viability caused by processing tissue with active type 2 immunity prevented direct measurement of IFNγ^+^ and IL17^+^ T cells in the distal SI of wild type *Tmu* colonized mice. Instead, we measured Th1 and Th17 induction in *Trpm5*^-/-^ mice, which lack the tuft cell taste chemosensory component TRPM5 and are insensitive to succinate-induced type 2 immunity^20,21^. Like *Tc* in wild type mice, *Tmu* stimulated expansion of IFNγ^+^, IL17^+^, and IFNγ-IL17 double positive CD4^+^ T cells in the distal SI of *Trpm5^-/-^* mice (Figure 3C and 3D). Altogether, these data suggest tritrichomonads can induce Th1 and Th17 immunity across the intestinal tract, but metabolic production of succinate by *Tmu* masks this response in the distal SI by driving type 2 immunity.

### Metabolic differences in tritrichomonads underly the divergent immune responses

Succinate excreted by *Tmu* stimulates the succinate receptor (GPR91) on tuft cells and drives type 2 immunity in the distal SI (Figure S4A)^20,21^. Because *Tc* does not stimulate type 2 immunity, we hypothesized that *Tc* does not produce succinate. We directly measured *in vivo* succinate concentrations in the ceca of *Tmu* and *Tc* colonized mice using liquid chromatography-mass spectrometry (LC-MS). *Tmu* colonization significantly increased the extracellular concentrations of cecal succinate compared to uncolonized controls, while *Tc* colonized mice showed no change in succinate levels (Figure 4A).

**Figure 4.**
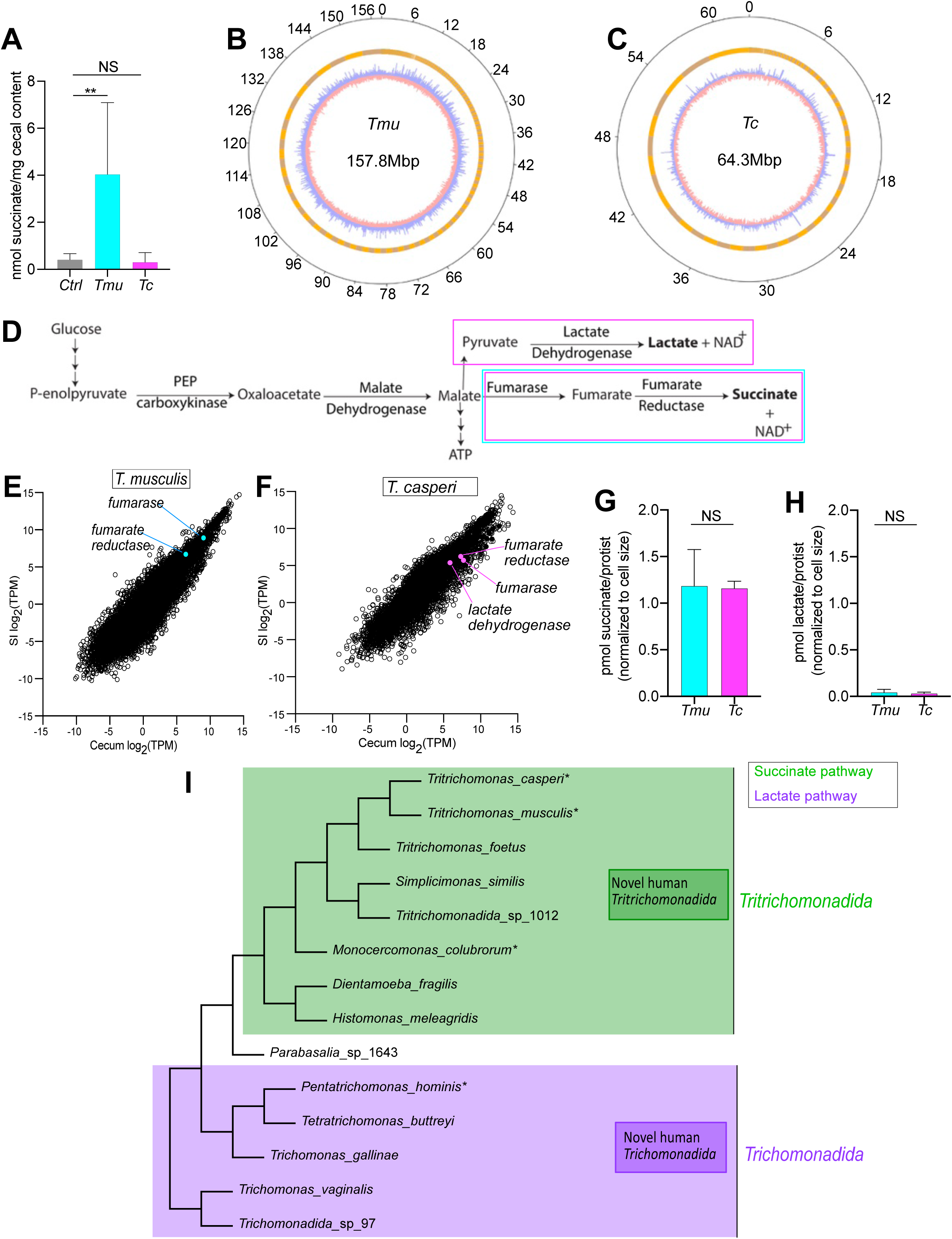
Metabolic differences in tritrichomonads underly divergent immune responses. (A) Extracellular succinate concentration in the cecal contents of uncolonized mice (*Ctrl*) or mice colonized with each protist for 3 weeks. (B, C) Genome assemblies of *Tmu* (B) and *Tc* (C). Outermost rings show position (in Mbp), middle ring shows genome assembly contigs, inner ring shows GC content in 10kb sliding windows (blue, above genome average; red, below genome average). Total genome size is at the center of each circle. (D) Schematic of metabolic pathways in parabasalid protists. Magenta boxes surround fermentative pathways found in *Tc*, cyan box surrounds the pathway found in *Tmu*. (E and F) Gene expression levels (in transcripts per million (TPM)) in the cecum vs. distal SI for *Tmu* (E) and *Tc* (F). Transcripts in the lactate and succinate fermentative pathways are labelled if present. (G and H) Intracellular succinate (G) and lactate (H) measurements from purified *Tmu* and *Tc* cell lysates. (I) Cladogram of parabasalids. Green shading shows predicted succinate producers. Purple shading shows predicted lactate producers. Asterisks signify new metabolic confirmations in this study. “Novel human *Tritrichomonadida/Trichomonadida*” represent novel human-associated parabasalids detected in Figure 1, but for which sequence information was not obtainable. Phylogenetic orders are labelled on the right. Graphs (A, G, H) depict mean with SD. *p<0.05, **p<0.01, NS, not significant by Student’s t test.

Two fermentative pathways are currently established in parabasalid protists based on studies of *T. vaginalis* and *T. foetus* metabolism^24,30^. *T. vaginalis* redox homeostasis converts pyruvate to lactate through the action of lactate dehydrogenase. In *T. foetus,* and presumably *Tmu*, redox balance occurs via conversion of malate to succinate by the sequential activity of fumarase and fumarate reductase. Lactate or succinate are then excreted as waste products from the protists. However, as with succinate, *Tc* colonization did not result in an increase in extracellular lactate in mouse cecal contents (Figure S4B).

To identify the metabolic pathways present in each protist, we sequenced the genomes of both *Tmu* and *Tc*. Massive numbers of repeat regions in parabasalid genomes make short-read based genome assemblies challenging^31–33^. Therefore, we used Nanopore long-read sequencing and short-read error correction to assemble these parabasalid genomes (Figure 4B, 4C, and S4C). As predicted, *Tmu* encodes the succinate production pathway, but not the lactate pathway (Figure 4D, cyan box). Despite the lack of succinate or lactate excretion by *Tc*, it encodes both the succinate and lactate production genes (Figure 4D, magenta boxes). To assess the relative contribution of each pathway to *Tc* metabolism, we performed RNA sequencing on *Tmu* and *Tc* isolated from the distal SI and ceca of mice. *Tmu* and *Tc* both expressed genes in the succinate pathway at high levels, whereas lactate dehydrogenase was less highly expressed in *Tc* (Figure 4E and 4F). Transcriptomics also confirmed that expression of these fermentative pathways does not vary drastically between the distal SI and cecum. Together, these data suggest that *Tmu* and *Tc* both use the succinate fermentation pathway, contrary to our hypothesis based on LC-MS measurements from cecal supernatants and the lack of type 2 immune induction by *Tc*.

To directly determine the usage of each fermentative pathway, we measured the intracellular concentrations of both metabolites in lysates from purified protists. Consistent with the genomic and transcriptomic analyses, cell extracts from purified *Tmu* and *Tc* contained high levels of succinate and low levels of lactate (Figure 4G and 4H). These intracellular succinate concentrations suggest that both protists use the succinate pathway to maintain redox homeostasis, yet cecal LC-MS measurements and the absence of tuft cell hyperplasia suggest *Tc* does not excrete this metabolite. Some bacteria can metabolize succinate into short chain fatty acids such as propionate and butyrate^34,35^. Therefore, we reasoned that *Tc* might also metabolize succinate in a similar manner. However, short chain fatty acids, including butyrate, propionate, and acetate, were not more abundant in lysates from *Tc* than *Tmu* (Fig. S4D). We next attempted to identify other potential succinate derivatives in *Tc* using additional mass spectrometry methods capable of identifying over 400 microbiota-produced metabolites^36^, but of the 48 metabolites we identified none had significantly higher abundance in *Tc* (Figure S4E). Altogether, these data indicate that although both protists use succinate production for redox homeostasis *Tc* does not excrete sufficient succinate into the intestinal lumen to stimulate type 2 immunity. However, the precise metabolic mechanism remains undefined.

We observed that *T. foetus*, *Tmu*, and *Tc* all use the succinate fermentative pathway and belong to the order *Tritrichomonadida*, whereas *T. vaginalis* uses the lactate pathway and belongs to the order *Trichomonadida*. This led us to hypothesize that an evolutionary split had occurred within the *Parabasalia* phylum, with lactate versus succinate production representing a branch point. To investigate this hypothesis, we analyzed the fermentative metabolism of two additional intestinal parabasalids: *P. hominis* (a human-associated *Trichomonadida*) and *Monocercomonas colubrorum* (a reptile-associated *Tritrichomonadida*). As predicted, *P. hominis* cultures contained high concentrations of lactate whereas *M. colubrorum* cultures contained high concentrations of succinate (Figure S4F). These data suggest that the phylogenetic classification of new and uncharacterized parabasalid species, such as those identified in this study, can help predict the production of the immunomodulatory metabolites lactate and succinate (Figure 4I). For example, this predictive framework suggests that the human-associated parabasalid *Trichomonadida_*sp_97 produces lactate, whereas the novel human-associated *Tritrichomonas* species likely use the succinate fermentative pathway (Figure 4I).

### *Tmu* and *Tc* occupy different nutritional niches within the microbiota

In addition to the microbiota’s metabolic output, the metabolic input of each species also shapes its interactions with other microbes and the host^37–39^. Plant polysaccharides such as dietary fibers are the most commonly utilized carbon source by the microbiota, and collectively are referred to as dietary Microbiota Accessible Carbohydrates (MACs)^40–42^. However, non-dietary MACs are also prevalent in the intestine. For example, mucus secreted by intestinal epithelial goblet cells provides a complex and glycan-rich nutrient source for microbes capable of breaking down its diverse glycosidic linkages^42–44^. To determine whether *Tmu* and *Tc* ferment dietary MACs, we altered the fiber content of mouse diets using defined chow similar to a previous study^21^. Fibers in these diets were restricted to inulin, a fermentable fiber that acts as a dietary MAC, and cellulose, a non-fermentable fiber that acts as a bulking agent and passes largely undigested through the intestine. Standard mouse chow, which contains a complex mixture of plant material, was used as a control (complex). *Tmu* abundance was high in mice fed complex chow and defined chow with inulin and cellulose (def-CI) or inulin alone (def-I) (Figures 5A and S5A). However, *Tmu* colonization was dramatically reduced when fiber was removed entirely (def-None) or cellulose was the sole fiber (def-C) (Figures 5A and S5A). In contrast, *Tc* colonization was completely independent of dietary MACs, as this protist’s abundance did not decrease on any of the defined diets (Figure 5B, S5B). These results suggest that *Tc* may rely on host-derived carbohydrates as a nutritional source, while *Tmu* colonization depends on dietary MACs.

**Figure 5.**
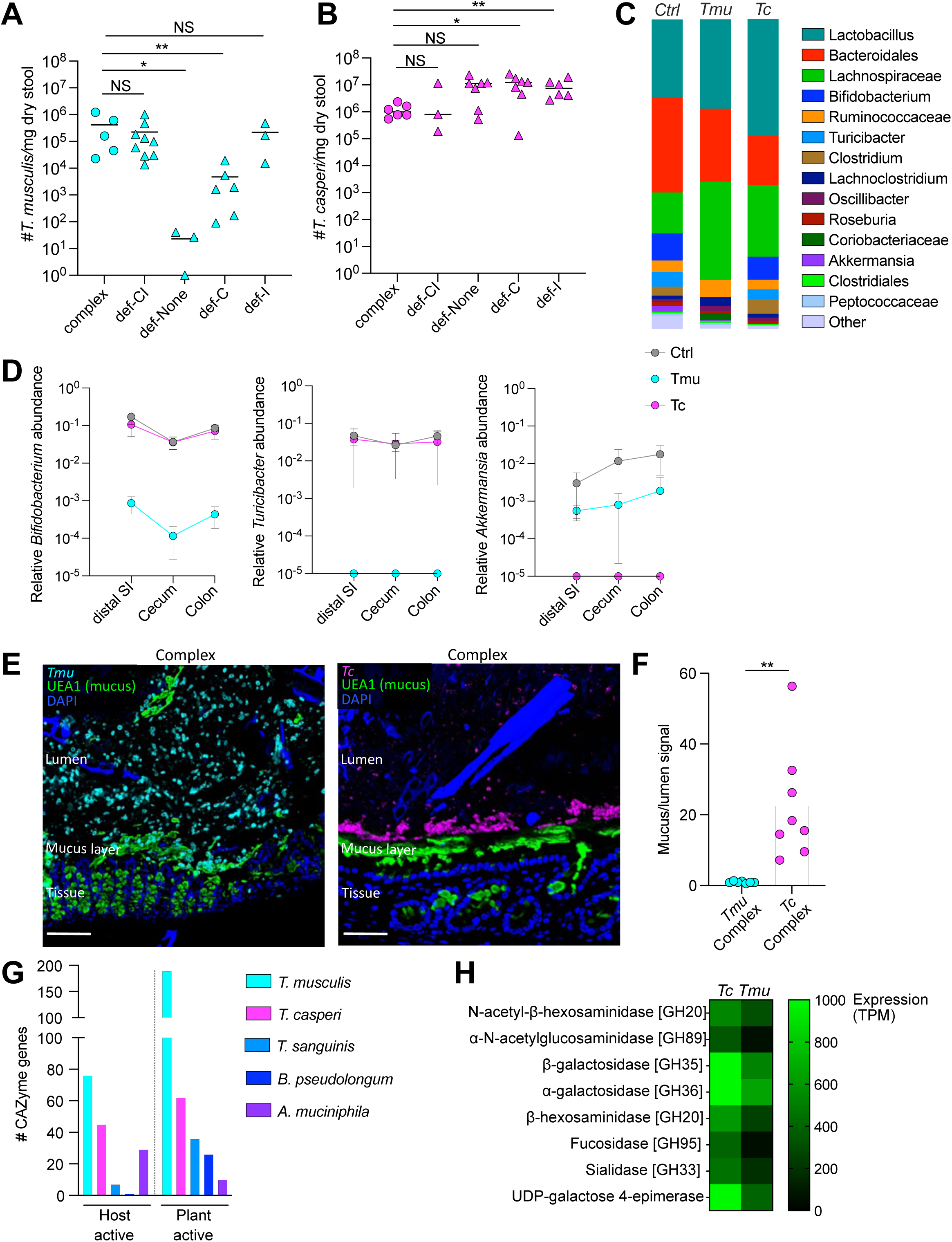
*Tmu* and *Tc* occupy different nutritional niches within the microbiota. (A) Abundance of *Tmu* and (B) *Tc* in the stool of mice fed diets with defined fiber compositions for two weeks or fed standard chow (complex) as a control. Fibers in the defined diets are limited to inulin (I), cellulose (C), or no fiber (None). (C) 16S rRNA amplicon sequencing in the colon of uncolonized (*Ctrl*) mice or mice colonized with *Tmu* or *Tc* for three weeks. The 15 most abundant taxa are shown, and data is the average abundance from five mice per group. (D) Relative abundance of bacteria depleted by *Tmu* and *Tc* colonization in each region of the intestine, from 16S Sequencing. (E) Representative microscopy showing localization of *Tmu* and *Tc* in the colons of mice fed complex chow. Scale bars are 100µm. (F) Quantification of protist localization by microscopy. Higher mucus/lumen protist signal indicates tighter localization to the colonic mucus layer. (G) Number of CAZymes encoded by protists and bacterial competitors with predicted activity against host and plant glycans. (H) Abundance (in TPM) of putative mucus-utilization genes in *Tc* and *Tmu* from transcriptomics data. CAZymes were identified by dbCAN2. For putative mucolytic CAZymes the glycoside hydrolase family (GH) is indicated in brackets. Graphs (A, B, D, F) depict mean, error bars depict SD in (D). Each symbol represents an individual mouse from one to three pooled experiments (A). **p<0.01, ***p<0.001, ****p<0.0001, NS, not significant by Mann-Whitney test (A, B) or Student’s t test (F).

The gut microbial ecosystem is a complex and species rich community, with most nutritional niches occupied by one or more community members competing for limited resources^37,45,46^. This scarcity of open niches can prevent new microbes from colonizing the gut and establishing themselves in the community, a phenomenon termed “colonization resistance”. However, commensal tritrichomonads such as *Tmu* and *Tc* are unusual in that they appear unaffected by colonization resistance. As few as one thousand *Tmu* or *Tc* protists can stably colonize SPF mice without niche-clearing manipulation. We reasoned that *Tmu* and *Tc* overcome colonization resistance due to one of two possibilities, (1) *Tmu* and *Tc* occupy unique and uninhabited niches within the gut ecosystem, or (2) *Tmu* and *Tc* outcompete other microbes that occupy their preferred niche. Due to the dearth of unoccupied niches, we favored the second hypothesis. Therefore, identifying microbes eliminated by tritrichomonad colonization could offer insights into the preferred nutritional niches of these protists.

To investigate trans-kingdom competition between tritrichomonads and bacteria, we performed 16S rRNA sequencing on the distal SI, cecum, and colon contents from mice with and without *Tmu* or *Tc* (Figures 5C, S5C, and S5D). In all three regions of the intestine, *Tmu* colonization dramatically reduced *Bifidobacterium pseudolongum* and *Turicibacter sanguinis* abundance (Figure 5C, 5D, S5C, and S5D). Alternatively, *Tc* colonization decreased *Akkermansia muciniphila* to undetectable levels (Figures 5C, 5D, and S5C-E). *Bifidobacteria* are prolific fiber degraders and *Turicibacter* abundance is positively correlated with fiber consumption^47,48^, consistent with *Tmu* outcompeting these bacteria for dietary MACs. *A. muciniphila* is a mucin specialist, suggesting that *Tc* may preferentially consume host mucus glycans.

In the colon, mucus forms a highly interlinked network that is tightly associated with the intestinal epithelium and acts as a barrier against microbial encroachment^2^. Mucolytic bacteria, such as *A. muciniphila*, preferentially colonize the mucus layer. To determine whether *Tc* also localizes to the colonic mucus layer, we used quantitative imaging to map the location of each protist along the mucosal-luminal axis of the intestine. *Tmu* was present at both the mucosal interface and in the lumen of the colon, consistent with a preference for dietary MACs (Figures 5E and 5F). *Tc* instead tightly co-localized with the colonic mucus layer, corroborating its preference for mucus glycans.

Mucus glycans and fiber are composed of diverse sugar structures that require a large repertoire of carbohydrate active enzymes (CAZymes) to catabolize. To investigate the glycan utilization potential of *Tmu* and *Tc*, we used dbCAN2 to identify CAZymes in the protist genomes^49^. Additionally, we annotated CAZymes in the genomes of the potential bacterial competitors *B. pseudolongum*, *T. sanguinis*, and *A. muciniphila.* Consistent with our hypothesis of nutritional trans-kingdom competition, *B. pseudolongum* and *T. sanguinis* were enriched for CAZymes active on plant polysaccharides, whereas *A. muciniphila* was enriched for host-active CAZymes (Figures 5G and S5F). However, *Tmu* and *Tc* each encoded a large number of both plant and host-active CAZymes, suggesting that these protists are capable of breaking down a wide variety of carbohydrate substrates. We therefore used the protist transcriptomes to determine the expression levels of mucus active CAZymes. *Tc* expresses CAZymes with predicted activity against all of the sugar subunits found on mucus glycans, and these CAZymes are members of Glycoside Hydrolase (GH) families associated with mucus degradation (Figure 5H)^27,43^. In particular, *Tc* expresses high levels of galactosidases, which can cleave the abundant galactose residues from mucus glycans. Galactose cannot be directly catabolized and must be converted to glucose to be oxidized for energy production. Consistent with a galactose-dependent metabolism, *Tc* also expresses high levels of UDP-galactose 4-epimerase used to convert galactose to glucose^50^. However, these putatively mucolytic CAZymes identified in *Tc* are also expressed by *Tmu,* albeit at lower levels of expression, suggesting that *Tmu* may also be capable of fermenting mucus glycans (Figure 5H).

### Dietary MAC deprivation causes *Tmu* to switch to mucolytic metabolism

Although *Tmu* colonization was markedly reduced when mice were fed the def-C diet, a small number of protists survived in a subset of these mice (Figure 5A and S5A). This observation, combined with the expression of mucolytic CAZymes, suggested that *Tmu* can utilize mucus glycans to survive dietary MAC deprivation. To investigate this possibility, we examined the localization of *Tmu* in the colons of mice fed cellulose as the sole fiber. The few remaining *Tmu* cells in def-C fed mice strongly co-localized with the mucus layer (Figure 6A and 6B).

**Figure 6.**
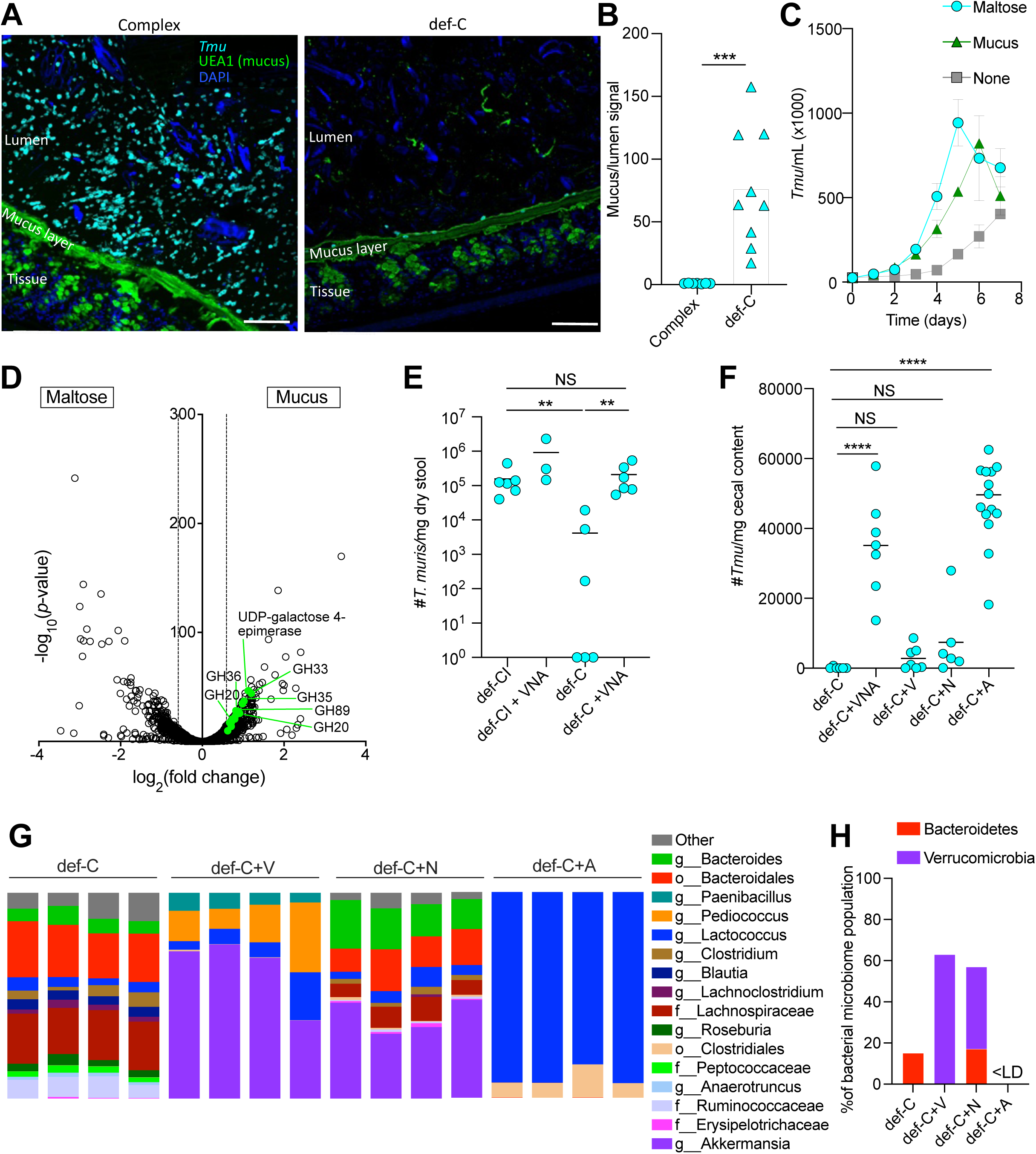
Dietary MAC deprivation causes *Tmu* to switch to a mucolytic metabolism. (A) Representative images of *Tmu* in the colons of mice fed complex chow (left) and defined chow with cellulose (def-C) chow (right) for two weeks. Scale bars are 100µm. (B) Quantification of microscopy on *Tmu* localization to the mucus layer when mice are fed complex chow or def-C chow. Higher mucus/lumen protist signal indicates tighter localization to the colonic mucus layer. (C) Growth curves of *Tmu* cultured *in vitro*, using the newly developed PBF medium, with maltose (cyan) or mucus (green) added as carbon sources, or with no defined carbon source added (grey). (D) Differential gene expression from *Tmu* grown *in vitro* with either maltose or mucus as a carbon source. Green data points indicate putative mucus-utilization genes with increased expression when grown in the presence of mucus. GH families for putative mucolytic CAZymes are labelled. Dashed lines delineate the fold change cutoff (1.5-fold). (E) *Tmu* abundance in the stool of mice fed def-CI or def-C chow for two weeks, with or without an antibiotic cocktail (VNA). Antibiotics used are Vancomycin, V; Neomycin, N; Ampicillin, A. (F) *Tmu* abundance in the cecal contents of mice fed cellulose chow for two weeks with different antibiotic treatments. (G) 16S rRNA sequencing of stool from *Tmu*-colonized mice fed def-C chow for two weeks with different antibiotic treatments. Each column represents an individual mouse. (H) Relative abundance of known mucolytic bacteria in mice from each group in (G). LD, limit of detection. Graphs (B, C, E, F, H) are plotted as mean, error bars depict SD in (C). Each symbol represents an individual mouse from one to three pooled experiments (B, E, and F). **p<0.01, ****p<0.0001, NS, not significant by Mann-Whitney test (E, F) or Student’s t test (B).

We next sought to study this metabolic shift in a reductionist model to understand the effects of specific substrates on protist growth and survival without the inherent complexity of the intestinal environment. Although some parabasalid species can be propagated *in vitro*, murine gut commensals such as *Tmu* and *Tc* do not grow with existing culture conditions. Although previous studies by our group and others have cultured these protists short-term to eliminate bacteria and fungi with antibiotics^15,19–21^, growth does not occur; instead, the protists slowly die over the course of several days. This method is sufficient to colonize mice with pure protists, but it does not allow for mechanistic investigation of protist biology. We created a new culture medium, which we named PBF, that supports the long-term culture of *Tmu* (Figure S6A). Despite the success of PBF for *Tmu* culture, it did not support the growth of *Tc*. Nevertheless, adding mucus to PBF medium improved *Tc* survival 5-fold, supporting the conclusion that this species ferments mucus glycans (Figure S6B).

Next, we cultured *Tmu* axenically with either the simple sugar maltose or with mucus as the primary carbon source and found that both glycans supported robust protist growth (Figure 6C). To confirm whether the CAZymes expressed by protists *in vivo* (Figure 5H) may support mucus glycan harvesting, we performed RNA sequencing on *Tmu* grown with either maltose or mucus (Figure 6D). *Tmu* grown with mucus significantly increased the expression of 16 CAZymes in GH families associated with mucolytic activity relative to protists grown with maltose. These upregulated CAZymes included the putative mucolytic genes identified above, supporting the hypothesis that both protists use these CAZymes to metabolize mucus glycans. In addition, the gene encoding the galactose-utilization enzyme UDP-galactose-4-epimerase was upregulated in mucus-grown *Tmu,* suggesting a metabolic shift towards fermenting mucosal sugars.

The discrepancy between *Tmu* survival on mucus *in vivo* (in which the protists barely survive) and *in vitro* (where the protists thrive) led us to consider what factors might account for this difference. One major distinction is the absence of bacteria in the *in vitro* cultures. To determine if mucolytic bacteria outcompete *Tmu* during dietary MAC deprivation *in vivo*, we treated *Tmu-*colonized mice with antibiotics before switching to the def-C diet. Strikingly, antibiotic treatment with a cocktail of vancomycin, neomycin, and ampicillin restored *Tmu* colonization in the absence of dietary MACs (Figure 6E). To narrow down the potential bacterial competitors, we tested the ability of each antibiotic to restore colonization. Neomycin and vancomycin were dispensable for *Tmu* survival, whereas ampicillin treatment completely restored colonization (Figure 6F and S6C). Consistent with ampicillin providing *Tmu* uncontested access to the mucus niche during fiber deprivation, the protists localized tightly to the mucus layer (Figure S6D and S6E).

To identify specific bacteria that outcompete *Tmu* during dietary MAC starvation, we profiled the changes to the bacterial microbiome during treatment with each of the individual antibiotics. Mice treated with ampicillin had undetectable levels of *Bacteroidetes* and *A. muciniphila*, known mucolytic members of the microbiota (Figure 6G, 6H, and S6F)^43^. In contrast, the mice treated with no antibiotics, vancomycin, or neomycin all had high levels of *Bacteroidetes*, *A. muciniphila*, or both. These results suggest that *Tmu* switches to a mucolytic metabolism during dietary MAC starvation but is largely outcompeted by the mucolytic bacteria *Bacteroidetes spp*. and *A. muciniphila*.

### Diet-protist interactions shape the host immune response

Next, we characterized dietary MAC fermentation by *Tmu* using axenic *in vitro* culture. Although inulin was sufficient for *Tmu* to colonize mice (Figure 5A), *Tmu* failed to grow in PBF medium with inulin as the carbon source (Figure 7A). To understand this apparent discrepancy, we investigated the localization of *Tmu* in mice fed def-CI chow, in which inulin is the sole dietary MAC. Similar to ampicillin treated mice starved of dietary MACs, protists were tightly associated with the mucus layer in mice fed def-CI chow (Figure 7B and 7C). This suggests that although inulin is sufficient for *Tmu* to colonize mice at high abundance, *Tmu* does not directly utilize inulin and instead ferments mucus glycans when inulin is the sole dietary MAC. Notably, many *Bacteroidetes* species can ferment both inulin and mucus, suggesting that these bacteria may utilize inulin when the fiber is present, thereby reducing competition with *Tmu* for mucus glycans^43,51^. *Bacteroidetes* abundance did not change in *Tmu*-colonized mice fed def-C chow compared to def-CI chow, whereas *Tmu* levels decreased in def-C chow (Figure 7D and 5A). This observation is consistent with *Bacteroidetes* outcompeting *Tmu* for mucus glycans when all dietary MACs, including inulin, are removed. However, this same dietary switch in *Tc*-colonized mice results in a significant decrease in *Bacteroidetes* abundance, consistent with the ability of *Tc* to outcompete mucolytic bacteria for the mucus niche (Figure 7D).

**Figure 7.**
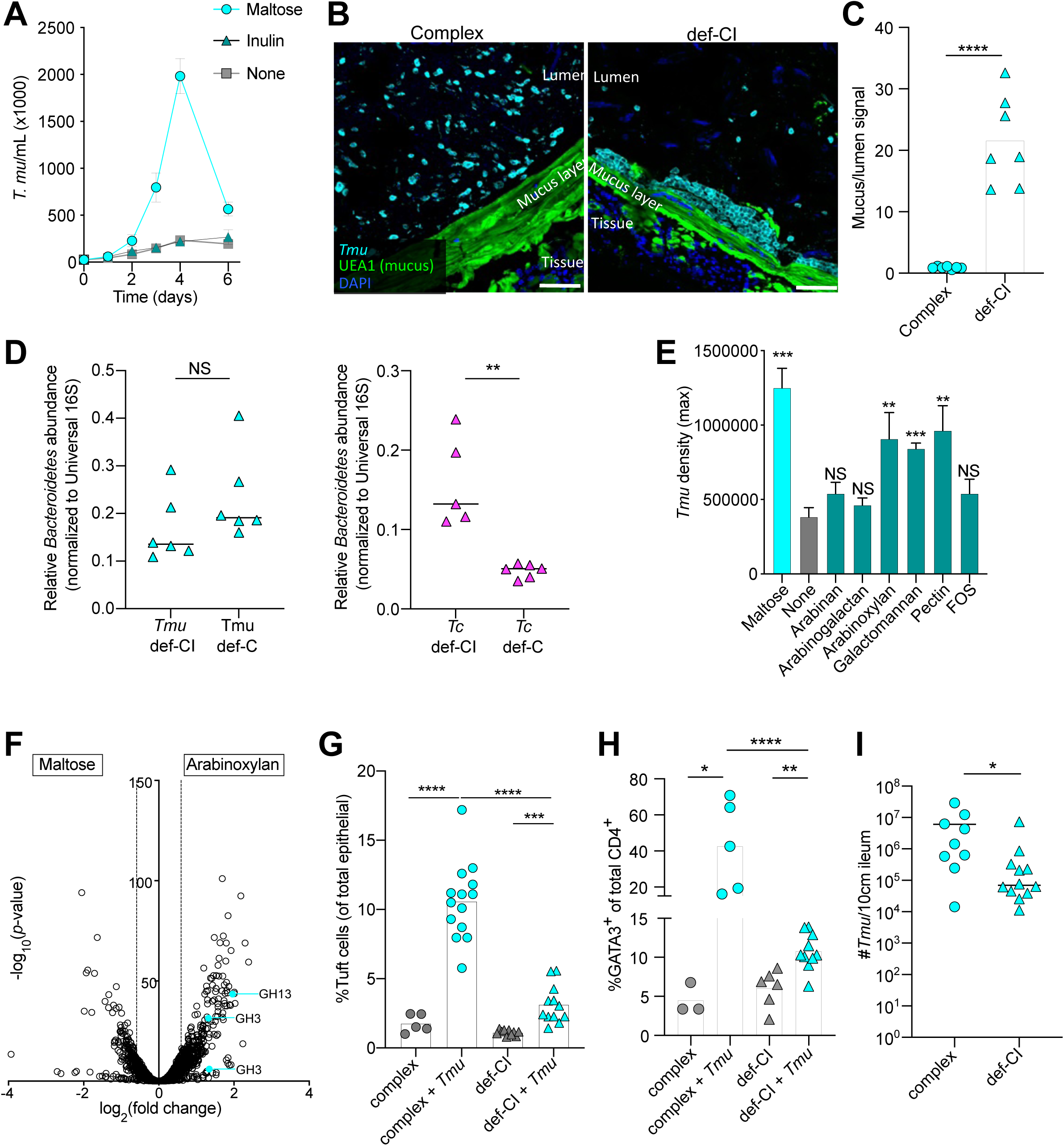
Diet-protist interactions shape the host immune response. (A) *In vitro* growth curve of *Tmu* cultured in PBF medium with maltose or inulin added as carbon sources, or with no defined carbon source added. (B) Representative images of *Tmu* in the colons of mice fed complex chow or defined chow with cellulose and inulin (def-CI) for three weeks. Scale bars are 50µm. (C) Quantification of microscopy on *Tmu* localization to the colonic mucus layer when mice are fed complex chow or def-CI chow. Higher mucus/lumen protist signal indicates tighter localization to the colonic mucus layer. (D) Relative abundance of *Bacteroidetes* bacteria in the stool of mice colonized by *Tmu* or *Tc*, with the fermentable fiber inulin or the non-fermentable fiber cellulose as the only fiber sources in the diet. (E) Maximum *Tmu* culture density when grown *in vitro* with different dietary MACs added as carbon sources to PBF medium. All statistical comparisons are relative to culture with no defined carbon source added. (F) Differential expression of *Tmu* transcripts grown *in vitro* with either maltose or arabinoxylan as a carbon source. Cyan data points represent upregulated CAZymes with potential activity towards arabinoxylan. Dashed lines delineate the fold change cutoff (1.5-fold). (G and H) Epithelial tuft cell (G) and lamina propria Th2 (H) frequency in the distal SI of mice colonized with *Tmu* or no protist (*Ctrl*) and fed complex or def-CI chow for 3 weeks. (I) *Tmu* abundance in the distal SI of mice fed complex chow and def-CI chow for 3 weeks. Data is plotted as mean with SD (A and E). Each symbol represents an individual mouse from two or three pooled experiments (C, D, G, H, and I). *p<0.05, **p<0.01, ***p<0.001, ****p<0.0001, NS, not significant by Mann-Whitney test (D, I) or Student’s t test (C, E, G, H).

To determine whether *Tmu* can ferment other dietary MACs, we tested a wider array of fermentable fibers as the carbon source in *Tmu* cultures. While *Tmu* failed to grow in maltose-free PBF medium supplemented with the fibers arabinan, arabinogalactan, and fructooligosacharides (FOS), the protist readily grew on arabinoxylan, galactomannan, and pectin (Figure 7E and S7A). To identify CAZymes responsible for fiber utilization by *Tmu*, we performed transcriptomics on *Tmu* cultured with either the simple sugar maltose or the fermentable fiber arabinoxylan (Figure 7F). We identified 3 CAZymes in GH families associated with fiber utilization that increased expression during growth on arabinoxylan, suggesting these CAZymes break down fiber polymers^27,43^.

Finally, we wondered whether diet and *Tmu* metabolic input would affect immune induction by the protist. When *Tmu*-colonized mice were fed def-C chow, and thus protists were largely depleted from the intestine, type 2 immune induction was completely ablated (Figure S7B and S7C). However, when *Tmu*-colonized mice were fed def-CI chow, and thus *Tmu* fermented mucus glycans, type 2 immune induction in the distal SI still occurred but was largely suppressed (Figure 7G and 7H). This suppression did not appear to be due to decreased succinate output by *Tmu* on a per-cell basis during mucus feeding, as *Tmu* grown on mucus *in vitro* showed no decrease in succinate excretion (Figure S7D). However, *Tmu* abundance in the distal SI of mice fed def-CI chow was substantially lower than in mice fed complex chow, suggesting that when fermenting mucus, this protist cannot colonize the SI at sufficiently high levels to induce type 2 immunity (Figure 7I). Altogether, these results demonstrate that alteration of dietary fiber intake can induce graded type 2 immune responses by *Tmu*.

## Discussion

The role of the eukaryotic microbiome, and commensal protists in particular, in shaping community structure and influencing host physiology is far less understood than their bacterial counterparts. This study reveals previously unrecognized phylogenetic diversity of commensal protists in the *Parabasalia* phylum in both mice and humans. Amongst the newly identified human-associated parabasalids are close relatives of the murine species, *Tmu* and *Tc*.

Although the species found in humans are distinct from the murine isolates, they reinforce the potential translational impact of functional studies using *Tmu* and *Tc*. We demonstrate numerous shared features between parabasalids, but importantly, our results also revealed that even minor phylogenetic differences can have meaningful biological consequences. Although *Tmu* and *Tc* are closely related species, they differ in the excretion of the fermentative waste product succinate, and they prefer distinct nutritional niches within the gut ecosystem. These metabolic differences lead to divergent small intestinal immune responses, distinct trans-kingdom interactions, and differential reliance on host dietary fiber. For example, succinate induced type 2 immunity in the SI has been a ubiquitous feature of previously characterized commensal tritrichomonads^19–21^, but *Tc* demonstrates that this is not universal. Additionally, all commensal parabasalids are assumed to ferment dietary fiber, but *Tmu* and *Tc* diverge in dietary fiber usage, leading to trans-kingdom competition with fiber or mucus digesting bacteria, respectively. However, comparing *Tmu* and *Tc* revealed unanticipated similarities as well. *Tc* increases Th1 and Th17 cells in the distal SI, whereas *Tmu* initiates type 2 immunity by stimulating tuft cells with succinate. However, *Tmu* colonization of mice lacking the tuft cell response to this metabolite instead exhibited a Th1 and Th17 response in the SI comparable to *Tc*. This result suggests both protists have the capacity to stimulate Th1 and Th17 immunity in the small intestine. Additional studies will be required to identify potential shared factor(s) that promote this immune response and determine the extent of its conservation in the *Parabasalia* phylum.

Our studies generated new tools and datasets for commensal tritrichomonads, which will eliminate critical barriers that have stymied mechanistic studies of these protists. The genomes of *Tmu* and *Tc* combined with metabolomic and transcriptomic datasets form a valuable foundation for future studies. Furthermore, the ability to grow these tritrichomonads in culture is an essential step for microbiological investigations. Previous studies, including our own, isolated tritrichomonads from mice, but the protists did not grow robustly in axenic *in vitro* culture^15,19–21^. This basic capability is particularly crucial for members of the microbiota because the complexity of their natural ecosystem confounds the ability to link phenotypes to single members of the community. In addition, the high degree of strain-level variation and potential for genetic drift make growth in culture and the ability to freeze/thaw isolates invaluable for increasing experimental rigor. Further, culture represents the first major hurdle for developing any microbe into a genetically tractable model.

Only two species of commensal parabasalid, *D. fragilis* and *P. hominis*, have been identified in the human gut. Previous studies also suggested industrialized microbiomes have decreased microeukaryotic prevalence^17,18^. Here, we identified multiple new human-associated commensal parabasalids, which were primarily found in the stool of individuals from non-industrialized populations. These new parabasalids further emphasize the need to consider the microbiome of diverse human populations and highlight the importance of understanding how different parabasalids influence human health. Our discovery that parabasalid production of the immunomodulatory metabolites succinate and lactate can be predicted based solely on phylogeny is a powerful tool. For example, this model predicts that *D. fragilis* and the newly discovered human-associated tritrichomonads produce succinate, and therefore may stimulate type 2 immunity in the small intestine. However, further work is required to refine this predictive model to account for alternative metabolic mechanisms, including those that may limit succinate excretion in protists like *Tc*.

## Limitations of the study

An important limitation of this study is that our identification of parabasalids in human microbiomes likely includes a high rate of false negativity. Despite the high sequencing depth in some of the metagenomic datasets, read depth was low for parabasalid ITS sequences in most samples. Thus, the prevalence of human-associated parabasalids identified here likely underestimates the true prevalence. Parabasalid-specific approaches will help to circumvent this issue in future studies. A second limitation of this study is that our predictive framework for fermentative pathways in parabasalids only allows for prediction of utilization of a pathway for redox homeostasis, and not excretion of the metabolite. Our discovery that *Tc* produces succinate but does not excrete sufficient quantities to stimulate type 2 immunity highlights the importance of this distinction. Identification of this metabolic mechanism will facilitate the prediction of excretion based solely on genomic data.

## Acknowledgements

We thank members of the Howitt lab, John Boothroyd, Markus Lakemeyer, Rebecca Gellman, and Fatima Enam for their helpful discussions and expertise. We also thank Dylan Maghini, Eli Moss, Roshni Patel, and Sandra Kong for their NGS expertise. We thank John Perrino and Ruth Yamawaki at the Stanford Cell Sciences Imaging Facility for expertise and help with sample processing and scanning electron microscopy. We thank Dr. R. Margolskee, Dr. J. von Moltke, and Amgen for kindly providing the *Trpm5^-/-^*, SMART13, and *Gpr91*^-/-^ mice, respectively. This work was supported by grants from the NIH (R01DK128292, K01DK113041, R01AI14862302, R01AI14375702, 5 T32 AI07290, and T32HL120824), a Jacob Churg award, and Stanford Pediatric IBD and Celiac Disease Research Program and the Stanford Maternal & Child Health Research Institute Seed Grant to M.R.H.. E.R.G. is supported by the Stanford Pediatric IBD and Celiac Disease Research Program and the Stanford Maternal & Child Health Research Institute.

## Author contributions

E.R.G, S.Z., P.T.W., M.M.C, M.R.O, A.S.B., and M.R.H. designed the experiments. E.R.G., S.Z., P.T.W., M.M.C., M.R.O., E.B.C., C.M.M., J.A.L., S.K.H. performed experiments and contributed intellectual expertise. J.L.S. and A.S.B., provided intellectual expertise. E.R.G. and M.R.H. conceived the study and wrote the manuscript, and all authors read and commented on the manuscript.

## STAR METHODS

### Key Resources Table

[See separate document]

## RESOURCE AVAILABILITY

### Lead contact

Further information and requests for resources and reagents may be directed to and will be fulfilled by the lead contact, Michael Howitt.

### Materials availability

Mouse and microbial strains used in this study will be made available upon request addressed to the lead contact.

### Data and code availability

Raw sequencing files and genome assemblies have been deposited at Stanford Digital Repository and are publicly available as of the date of publication (links to each dataset can be found in Key Resources Table). Any additional information required to reanalyze the data reported in this paper is available from the lead contact upon request.

### Animals

C57BL/6 WT and *Caspase1*^-/-^ mice were purchased from Jackson Laboratory. Mice from Jackson Laboratory are protist-negative, as verified by PCR on the stool and examination of cecal contents. *Tprm5*^-/-^ and *Gpr91^-/-^* mice were generously provided by Dr. Robert Margolskee and Amgen, respectively, under materials transfer agreements. The IL13 reporter mouse line (SMART13 mice)^54^ were generously provided by Dr. Jakob von Moltke. All mice were used at 6-12 weeks of age. Experiments were performed using age and gender matched groups. All animal procedures used in this study were approved by the Institutional Animal Care and Use Committee (IACUC) at Stanford School of Medicine.

All mice were housed in individually ventilated cage systems and maintained at specific pathogen free (SPF) health status, unless stated otherwise. Colonies of protist-negative and protist-colonized mice were maintained and handled separately. Protist colonization status was routinely verified by PCR on stool and examination of cecal contents by light microscopy. Cage bottoms were covered with autoclaved bedding and enrichment material. Mice were handled within HEPA filtered air exchange stations, and all surfaces and tools were sanitized using Virkon^TM^ S disinfectant before and after handling. All cage changes were performed by lab staff, who were required to wear personal protection equipment (lab coats, gloves, disposable sleeves) to handle mice used in this study.

Custom diets for *in vivo* fiber experiments were obtained from Research Diets, Inc. The diets were the same as those used previously^21^, with diet IDs D11112201 (Defined chow with cellulose and inulin), D11112229 (no fiber), D17030102 (cellulose), D11112226 (inulin). For antibiotic treatment experiments, mice were given antibiotics in the drinking water, which was changed every 3 days. Antibiotics and concentrations used were: vancomycin (0.5g/mL), neomycin (1g/mL), ampicillin (1g/mL). For succinate treatment, disodium succinate (100mM) was added to the drinking water and changed every 3 days.

### Method details

#### Identification of protists in mouse stool

Mice at Stanford University animal facilities were screened for commensal protists by extracting DNA from stool using the DNeasy Powersoil Pro Kit (Qiagen) according to manufacturer instructions. Samples were then used as input for PCR using pan-parabasalid primers (forward: 5’-CCACGGGTAGCAGCA-3’, reverse: 5’-GGCAGGGACGTATTCAA-3’). These primers generate an approximately 1.1kb amplicon of the parabasalid internally transcribed spacer (ITS) sequence, which was sequenced by Sanger Sequencing for phylogenetic classification. Phylograms were generated using MAFFT^55^.

#### Isolation of *Tritrichomonas spp.* from mice

Protists were purified as described previously^19^, with modifications. Briefly, cecal contents were harvested from WT C57BL/6 mice and passed over a 40µm strainer. Contents were then washed three times in PBS and purified at the interface of a 40%/80% Percoll (GE healthcare) gradient, after centrifugation at 1000xg for 10minutes with no brake. The number of viable protists was quantified using a hemocytometer, and 1x10^6^ trophozoites were inoculated per mL of PBF growth medium. The PBF medium was supplemented with 1x Penicillin/Streptomycin (Caisson Labs), 2.5µg/mL Amphotericin B (Sigma Aldrich), 100µg/mL Vancomycin (Alfa Aesar), 12.5µg/mL Chloramphenicol (Sigma). Protists were then cultured overnight in an anaerobic chamber (Coy Labs) at 37°C.

#### Colonization of mice with protists

Mice were colonized with commensal para15.basalid protists as described previously^19^ with slight modifications. Briefly, protists were resuspended in PBS and mice were colonized with 1x10^5^ protists per mouse by oral feeding. Colonization was verified after 7 days by qPCR on the stool using *Tmu* or *Tc* specific primers. For *Tc* colonization of mice, protists were isolated directly from mouse ceca one day prior to colonization. For *Tmu* colonization of mice, protists were cultured *in vitro* for no more than 20 passages prior to colonization of mice.

#### Scanning Electron Microscopy (SEM)

SEM was performed on *Tmu* and *Tc* as described previously^19^. Briefly, protists were isolated from the ceca of SPF mice as described above and adhered to poly-lysine coated coverslips. Samples were then fixed in 2.5% glutaraldehyde in 0.1M cacodylate pH 7.2. Samples were then rinsed and post-fixed for 30min in 1% OsO_4_ in 0.1M cacodylate buffer and dehydrated in a graded series of ethanol prior to critical point drying with liquid CO_2_. Protists were then sputter-coated with 5nm platinum and examined with a Zeiss Sigma scanning electron microscope.

#### Collating previously published human gut metagenomic samples

Prevalence of parabasalid species across lifestyle was characterized using a curated collection of 1,800 metagenomes including samples from “industrial”, “transitional” and “hunter-gatherer” populations that were reported previously^28^. Due to the sensitivity of this analysis to read depth, we removed studies that had a median read depth lower than 5 giga base pairs.

#### Discovery of putative novel ITS sequences

We compiled a set of Parabasalid internal transcribed spacer (ITS) sequences derived in this study or from public databases. We used these sequences as “bait” to mine for novel ITS sequences in the contigs of metagenome assemblies of samples derived from Tanzania, Nepal or California^28^. The comparison between bait and assembly contigs was performed by mashmap^56^. A phylogenetic tree of these putative novel ITS sequences, along with our bait ITS sequences, was created using FastTree^57^.

#### Species prevalence analysis

All reads were mapped against a database of Parabasalid ITS sequences derived in this study or from public databases (Bowtie2)^58^. Resulting mappings were processed using inStrain profile (v1.2.14)^59^ and CoverM v0.4.0 (https://github.com/wwood/CoverM). Species with more than one read aligning to the ITS sequence at a breadth greater than 20% were considered present and prevalence was calculated as the percentage of metagenomes in which the species was present.

#### High molecular weight DNA extraction and Nanopore Sequencing

To obtain pure protist DNA for genome sequencing, *Tc*-colonized mice were treated with an antibiotic and antifungal cocktail in the drinking water (0.5g/L Vancomycin, 1g/L Neomycin, 1g/L Ampicillin, 0.2g/L Amphotericin B) for 6 days. *Tc* was then purified from the cecal contents as described above, with the modification of two rounds of Percoll purification, instead of one. For *Tmu*, protists were instead axenically cultured. High molecular weight DNA from each protist was then isolated by gently resuspending pelleted protists in lysis buffer (PBS+0.5% sodium dodecyl sulfate), using a wide-bore pipet tip. Lysed cells were transferred to DNA LoBind tubes (Eppendorf) and incubated with Monarch RNase A (NEB) for 2 hours at 37°C. Next, Proteinase K (Qiagen) was added, and samples were incubated for 2 hours at 50°C. Nucleic acid was then isolated using two rounds of extraction with Phenol:Chloroform:Isoamyl alcohol (Sigma). To avoid pipetting during transfer steps, vacuum grease (Dow Corning) was added as a phase-lock gel. Protein contaminants were then precipitated using 2M Ammonium Acetate, and then DNA was precipitated overnight in ethanol. Next, DNA was pelleted by centrifugation at 8000xg, and DNA pellets were washed three times with 70% ethanol. DNA was then resuspended in water and used for library preparation and sequencing using the Nanopore Ligation Sequencing Kit and Minion Flow Cell (Oxford Nanopore).

#### Genome assembly and annotation

Genome assemblies for both *T. casperi* and *T. musculis* were constructed using Flye^60^, v. 2.9-b1768 with default parameters. Short read libraries were then mapped against the genome assemblies with Bowtie2^58^, v. 2.4.4 and resulting alignments were used for assembly polishing with pilon^61^. Genes were predicted with MetaEuk^62^, v. 5.34c21f2 with Uniref90 as the reference protein dataset. Genome completeness was assessed with BUSCO^63^, v5.0.0 in protein mode with Eukaryota ODB10 lineage marker gene set. Genome diagrams were generated using DNAPlotter^64^, v. 18.2.0.

#### RNA-Sequencing

RNA-Sequencing was performed on *Tmu* and *Tc in vivo* by first isolating the luminal contents from the distal small intestine and cecum of 3 mice per group. Contents were strained over 40µm filters, then washed with ice-cold PBS to remove debris. To minimize potential for sample processing to affect transcriptomes, samples were then immediately resuspended in Qiazol (Qiagen). RNA was purified using the Direct-zol RNA Miniprep kit (Zymo Research). RNA-Seq libraries were then generated using the KAPA Stranded RNA-Seq kit with poly-dT enrichment of mRNA transcripts, and libraries were sequenced on an Illumina HiSeq instrument.

Transcriptomics on the *in vitro* grown *Tmu* samples was performed on protists grown for 3 days in the condition of interest. Cultures (3 biological replicates per condition) were pelleted at 700xg for 5min at 4°, then resuspended in Qiazol. RNA was then isolated using the Direct-zol RNA Miniprep kit (Zymo Research), as above. Library preparation and sequencing was performed at Azenta Life Sciences (South Plainfield, NJ). Libraries were constructed using the TruSeq Stranded mRNA kit (Illumina), with poly-dT enrichment of mRNA transcripts. Samples were sequenced on a HiSeq instrument (Illumina).

Analysis of all transcriptomics datasets was performed using alignment-based analysis, using the newly sequenced *Tmu* and *Tc* genomes. Briefly, Illumina adapters were trimmed and low-quality sequence was removed using Trimmomatic^65^. Reads were then aligned to the respective genome using Burrows Wheeler Aligner^66^. Samtools^67^ was used to convert corresponding files to binary alignment and map (bam) files and sort the reads by position. Next, featureCounts^68^ was used to quantify read depth of each genome feature. Differential expression analysis was performed using DESeq2^69^.

#### Protist enumeration

Enumeration of protists from the small intestinal tissue was performed as described previously^19^, with modifications. Briefly, the distal 10cm of small intestine was removed and flushed with ice-cold sterile PBS using a 19-gauge feeding needle. The contents were then pelleted and the supernatant was aspirated. Genomic DNA was isolated using the Powersoil Pro Kit (Qiagen) according to manufacturer instructions. To detect and enumerate protists, quantitative PCR (qPCR) was performed using PowerUp SYBR Green Master Mix (Applied Biosystems). For *T. musculis*, the following primers specific to the 28S rRNA gene were used: 5’-GCTTTTGCAAGCTAGGTCCC-3’and 5’- TTTCTGATGGGGCGTACCAC-3’^19^. For *T. casperi* detection, primers were designed that specifically recognized this protist: 5’- AGGTTACTGAATCATACATGCGT-3’ and 5’-GCAGGAGTTGCTTTCATTGTG-3’. qPCRs were run on a QuantStudio 3 Real Time PCR instrument (Thermo Fisher). To convert qPCR values into protist numbers, protists were isolated and counted using a hemocytometer before extracting genomic DNA and analyzing a dilution series by qPCR. These results were used to create a standard curve, and regression analysis was used to convert Ct values to protist numbers, as has been described previously^15,19^.

Quantification of protists in the stool was performed similarly. Stool pellets were collected from mice and weighed. A fraction of the stool was dried and used to calculate the water content of the stool, while the rest was used for DNA extraction. Quantification was performed using the same qPCR primers as above. Protists were quantified in the cecum by weighing the cecal luminal contents, then resuspending in PBS. Live trophozoites were then counted using a hemocytometer and normalized to cecal weight.

#### Isolation of Epithelial Cells for Flow Cytometry

Ileal epithelial cells were isolated as described previously^19^. Briefly, the terminal 10cm of the small intestine was removed and trimmed of fat. Luminal contents were gently flushed with 10mL of ice-cold PBS, then the tissue was opened longitudinally. Peyer’s Patches were removed and the tissue was incubated on ice in PBS, 5mM HEPES pH 7.4, 2% heat-inactivated fetal bovine serum (iFBS), 1mM DTT. Tissue was then transferred to pre-warmed PBS, 5mM HEPES, 2% iFBS, 5mM EDTA and shaken at 200rpm for 15 minutes at 37°, followed by vigorous shaking. This was repeated and epithelial cells from both fractions were combined and washed with PBS. Tissue was then digested in DMEM containing 10% iFBS, 0.5U/mL Dispase II (StemCell Technologies), 50µg/mL DNase (Roche) for 10 minutes at 37°C. The cells were then filtered over 40µm strainers and washed with PBS, 2% iFBS, 1mM EDTA. Cells were then incubated for 10 minutes with anti-CD16/CD32 (clone 93, Biolegend) and then stained with the following antibodies: PacBlue-conjugated anti-CD45 (clone 30-F11, Biolegend), PE/Cy7-conjugated anti-EpCam (clone G8.8, Biolegend), and Alex647-conjugated anti-SiglecF (clone E50-2440, BD Biosciences). Cell viability was assessed by propidium iodide staining (Biolegend). Cells were then analyzed on a BD FACSCanto II.

#### Lamina propria cell isolation and flow cytometry

The distal 10cm of small intestine, or the entire colon, were collected and epithelial cells were removed as described above. The remaining tissue was then minced and digested in RPMI 10% iFBS, 0.1 units/mL Dispase II (StemCell Technologies), 50µg/mL DNase (Roche), collagenase A (Roche) (0.25mg/mL for ilea, 0.5mg/mL for colons). Tissue was digested for 30 minutes followed by a second digestion for 40 minutes. Digested tissue was passed through a 40µm strainer, and the cells were counted using a Cellometer Auto T4 (Nexcelom Bioscience). For cytokine measurements, cells were stimulated in RPMI supplemented with 10% iFBS, Pen/Strep, PMA, Ionomycin, and GolgiPlug. Cells were incubated at 37° for 4 hours to allow stimulation to occur before being harvested. Cells were then incubated with anti-CD16/32 (clone 92, BioLegend) before surface staining. For T cell analysis, cells were stained with PacBlue-conjugated anti-CD45 (clone 30-F11, BioLegend), FITC-conjugated anti-CD4 (clone GK1.5, BioLegend), and PerCP/Cy5.5-conjugated anti-CD3 (clone 17A2, BioLegend). For ILC2 analysis, cells were stained with PacBlue-conjugated anti-CD45 (clone 30-F11, BioLegend), PerCP/Cy5.5-conjugated anti-CD127 (clone A7R34, BioLegend), PE/Cy7-conjugated anti-KLRG1 (clone 2F1, BioLegend), PE-conjugated anti-human CD4/Smart13 (clone RPA-T4, BioLegend), and a custom FITC conjugated lineage cocktail of antibodies against NK1.1 (clone PK136, BioLegend), CD19 (clone 6D5, BioLegend), CD4 (clone GK1.5, BioLegend), CD8 (clone 53-6.7, BioLegend), CD64 (clone S18017D, BioLegend), CD11b (clone M1/70, BioLegend), and TER119 (clone TER-119, BioLegend). Cells were then stained with the viability dye Zombie-NIR (BioLegend) according to manufacturer instructions prior to fixation with the Foxp3 Transcription factor staining buffer set (eBioscience). For T cell analysis, fixed cells were permeabilized and stained with PE-conjugated anti-IL17 (clone 9B10, Biolegend) and APC-conjugated anti-IFNγ (clone XMG1.2, eBioscience), or with Alexa647-conjugated anti-GATA3 (clone L50 823, BD) or Alexa647-conjugated anti-FOXP3 (clone MF-14, BioLegend). Cells were then analyzed on a BD FACSCanto II.

#### Tissue preparation and staining for immunofluorescence

Tissue sections of interest were removed and fixed in methacarn fixative (Fisher Scientific) for 48 hours. Tissue was then washed three times in 100% ethanol, followed by three washes in Histo-Clear II (Electron Microscopy Sciences). Fixed tissue was embedded in paraffin wax prior to sectioning (4µm) using a microtome. For immunofluorescence, tissue was first deparaffinized as has been described previously^70^, then blocked with blocking buffer (PBS supplemented with 3% BSA, 0.1% Saponin, 2% goat serum). Sections were then stained with the anti-*Tmu* antibody (generated by Rockland Scientific) and anti-E Cadherin (clone 36/E) overnight. After 3 washes in PBS, sections were stained with secondary antibodies fluorescently conjugated to Alexa488 (*Tmu*) and Alexa660 (E Cadherin). DAPI (Sigma Aldrich) and rhodamine-conjugated Ulex Europaeus Agglutinin I (Vector Laboratories) were added during the secondary staining step as well. After 3 PBS washes, tissue was mounted with Vectashield (Vector Labs) and imaged on a Zeiss LSM 700 confocal microscope.

#### Microscopy quantification of protist localization to the colonic mucus layer

One to three images were taken for each mouse for quantification. A 40µm tall mucosal region of interest (ROI) was drawn at the luminal edge of the inner colonic mucus layer. An ROI of identical size was placed in a representative region of the lumen, a minimum of 100µm away from the mucus layer. Fluorescence signal from the protist channel was measured, and the ratio of the signal from the mucosal ROI to the luminal ROI was calculated. The average of the ratios from each image per mouse was then calculated as the mucus/lumen signal.

#### Metabolite measurements

For intracellular metabolite analysis, mice were treated with an antibiotic cocktail in the drinking water (0.5g/L Vancomycin, 1g/L Neomycin, 1g/L Ampicillin, 0.2g/L Amphotericin B) for 6 days. Protists were then isolated as described above and lysed in 1mL 80% methanol. Cellular debris was removed through centrifugation, followed by filtration using 0.22µm filters, snap frozen in liquid nitrogen, and stored at -80°. Prior to liquid chromatography mass spectrometry (LC/MS) analysis, intracellular extracts (800 µL) were dried in a vacuum concentrator at room temperature for approximately 4 hours then resuspended in 50 µL of HPLC-grade water spiked with ^13^C-labeled metabolites as an internal standard.

For metabolite measurements from cecal supernatants, cecal contents were squeezed into tubes and weighed, prior to gentle resuspension in sterile saline solution (0.15M NaCl). Samples were then pelleted at 500xg for 5min at 4°, and supernatants were transferred to clean tubes. Samples were then pelleted at 2000xg for 10min at 4°, and then supernatants were filtered over 0.22µm filters, snap frozen in liquid nitrogen, and stored at -80°. Prior to LC/MS analysis, cecal extracts were diluted in HPLC-grade water spiked with ^13^C-labeled metabolites as an internal standard.

For organic acid measurements, samples (5 µL) were run in reverse phase on a 2.1 mm x 150 mm Waters Acquity UPLC BEH C18 column with 1.7 µm packing. The column was fitted with a Vanguard pre-column of the same composition. Analytes were separated on an Agilent 1290 Infinity II UPLC (binary pumps) and detected using an Agilent 6545 LC/MS Quadrupole Time-of-Flight (qTOF) instrument equipped with a dual jet stream electrospray ionization source (ESI) operating under extended dynamic range (EDR 1700 m/z) in the negative (ESI-) ionization mode. The mobile phase solvents consisted of 20 mM ammonium formate pH 2.9 (A) and acetonitrile (B). The separation began with 99.5% A and 0.5% B for 4 min at 0.4 mL/min, then B was increased to 99.5% at 0.5 mL/min and held for 1.5 min, at which time the flow rate was increased to 0.8 mL/min. At 11.5 min, solvent B was reduced to 0.5% and the flow rate was reduced to 0.5 mL/min for 2.5 min, at which time it was further reduced to 0.4 mL/min. The separation ended at 14 min. Specific ion source parameters included: Drying gas temperature was 150°C at 12L/min, sheath gas temperature was 325°C at 12 L/min, Nebulizer was at 45 psig, capillary voltage was 2000 V, nozzle voltage was 0V, fragmentor voltage was 90 V, skimmer voltage was at 35 V, octopole RF was 750V. For quantitation purposes, standard curves of all analytes were prepared in HPLC water spiked with ^13^C-labeled metabolites as an internal standard. Metabolite levels were determined by manually integrating the area under each chromatographic peak and their concentrations in the samples determined by interpolating the peak area for each compound on the corresponding standard curve.

For short chain fatty acid measurements, a LC-MS-based SCFA/OA quantification method was adapted^71^. Briefly, cell pellets diluted in extraction buffer containing: 80% HPLC-grade water (Fisher), 20% HPLC-grade acetonitrile (ACN; Fisher), and labeled isotopes of each SCFA/OA measured (2.5 μM d3-acetic acid [Sigma-Aldrich], 1 μM propionic-3,3,3-d3 acid [CDN Isotopes], and 0.5 μM butyric-4,4,4-d3 acid [CDN Isotopes]). Acid-washed beads were added to the samples and the samples were subsequently shaken at 30 Hz for 10 minutes. The supernatants were then derivatized in a combination of 200 mM 3-nitrophenylhydrazine hydrochloride (Sigma-Aldrich; dissolved in 50% ACN and 50% water) and 120 mM 1-ethyl-3-(3- dimethylaminopropyl)carbodiimide hydrochloride (Pierce; dissolved in 47% ACN, 47% water, and 6% HPLC-grade pyridine [Sigma-Aldrich]). The reaction was incubated at 37°C for 30 min and subsequently diluted 50X prior to mass spectrometry analysis.

Analyses were carried out using an Agilent 6470 triple-quadrupole LC-MS and a Waters Acquity ultraperformance liquid chromatography (UPLC) ethylene-bridged hybrid (BEH) C18 column (100-mm length, 2.1-mm inner diameter, 130-Å pore size, 1.7-μm particle size). The mobile phases were water-formic acid (100:0.01, vol/vol; solvent A) and acetonitrile-formic acid (100:0.01, vol/vol; solvent B). Quantification of analytes was done by standard isotope dilution protocols. In brief, serial dilutions of a SCFA/OA standard solution (10 mM, 1 mM, 0.1 mM, 0.01 mM, 0.001 mM, and 0 mM) were derivatized as described above and included in each run to verify that sample concentrations were within linear ranges. For samples within linear range, analyte concentration was calculated as the product of the paired internal standard concentration and the ratio of analyte peak area to internal standard peak area.

Untargeted LC-MS metabolomics was performed as described previously^36^ using C18^+^ and C18^-^ modes.

#### Dietary fiber experiments

For experiments involving manipulation of dietary fiber, mice (fed normal complex chow) were colonized by 1x10^5^ protists between 4-6 weeks of age. Ten days later, the diets were changed to chows with defined fiber compositions. Stool was collected for protist quantification two weeks later, and mice were euthanized for collection of tissue for additional analyses. For experiments involving manipulation of dietary fiber and antibiotic treatment, mice were colonized by protists as above. Ten days later, mice were started on the indicated antibiotic treatment, followed by diet switch three days after the start of antibiotics.

#### 16S Bacterial Profiling

For 16S bacterial profiling, mice were colonized by 1x10^5^ protists between 4-6 weeks of age, and stool was collected 3 weeks later for microbiome analysis. DNA was extracted using the Powersoil Pro Kit (Qiagen). The DNA samples were prepared for targeted sequencing at Zymo Research (Irvine, CA). Libraries were made using the *Quick*-16S NGS Library Prep Kit using Primer Set V3-V4 (Zymo Research, Irvine, CA). Samples were sequenced on an Illumina MiSeq with a v3 reagent kit. Unique amplicon sequences were inferred from raw reads using the Dada2 pipeline^72^. Chimeric sequences were removed, and taxonomy assignment was performed using Uclust from Qiime2^73^. Taxonomy was assigned with the Zymo Research 16S Database. Alpha-diversity and composition analyses were performed with Qiime.

Absolute abundance quantification was performed using qPCR. A standard curve was made with plasmid DNA containing one copy of the 16S gene, prepared in 10-fold serial dilutions. The primers used were the same as those used in targeted library preparation. The equation generated by the plasmid DNA standard curve was used to calculate the number of gene copies in the reaction for each sample. The PCR input volume was used to calculate the number of gene copies in each DNA sample. The number of genome copies per microliter of DNA in each sample was calculated by dividing the gene copy number by an assumed number of gene copies per genome. The value used for 16S copies is 4. The amount of DNA per microliter of sample was calculated using an assumed genome size of 4.64x10^6^bp, the genome size of *Escherichia coli*. The mass of the stool sample used for DNA extraction was then used to calculate the number of genome copies per milligram of stool.

#### Bacterial relative abundance measurements by qPCR

*Akkermansia muciniphila* and *Bacteroidetes* quantification was performed using DNA isolated from stool as described above. Relative abundance of bacterial groups were then quantified using the following primers: *Akkermansia muciniphila* 5’-CAGCACGTGAAGGTGGGGAC-3’, 5’- CCTTGCGGTTGGCTTCAGAT-3’^74^. *Bacteroidetes* 5’-GTTTAATTCGATGATACGCGAG-3’, 5’- TTAASCCGACACCTCACGG-3’^75^. Universal 16S 5’-AAACTCAAAKGAATTGACGG-3’, 5’- CTCACRRCACGAGCTGAC-3’^75^. qPCRs were performed using PowerUp SYBR Green Master Mix (Applied Biosystems) on a QuantStudio 3 Real-Time PCR machine (Thermo Fisher). Ct values for bacterial groups of interest were then normalized to total 16S for relative abundance analysis.

#### Quantification and Statistical Analysis

Groups were compared using Prism v9 software (GraphPad) using the two-tailed unpaired Student’s t test or Mann-Whitney test where appropriate. Differences were considered statistically significant when p::0.05.

## Supplemental figure legends

**Figure S1. Discovery of novel human- and mouse-associated parabasalids, related to Figure 1**.

(A) Length of *Tmu* and *Tc* trophozoites based on SEM images. (B) Alignment of partial ITS sequences from *Tmu* and *Tc*. (C) *Tc* abundance in the stool of mice colonized for 3 weeks or 3 months. Data is plotted as mean and each symbol represents an individual mouse. ****p<0.0001, NS, not significant by Students T test. (D) Average Nucleotide Identity (ANI) of the parabasalid reads in human metagenomic samples from (E). Imperfect ANI to the specified reference sequence indicates a relative of the reference protist. First author of the cohort study, country of origin, and population type (“Industrial”, “Transitional”, Hunter-Gatherer (“H-G”) is indicated. Columns and rows are hierarchically clustered based on Euclidean distance. Cells corresponding to protists not identified in a population are colored grey. (E) Prevalence of parabasalids in human metagenomics datasets using mapping-based analysis, without exclusion criteria based on sequencing depth. Parabasalids with imperfect matches in (D) represent relatives of the reference protist species. (F) Percentage prevalence of *D. fragilis* as a function of sequencing depth, separated by industrialization status. (Abbreviations: TZA=Tanzania, NEP=Nepal, MNG=Mongolia, USA=United States of America, MDG=Madagascar, KAZ=Kazakhstan, SWE=Sweden, HMP=Human Microbiome Project, IND=Indonesia, ISR=Israel, FJI=Fiji, ETH=Ethiopia, THA=Thailand, PER=Peru, CMR=Cameroon).

**Figure S2. *Tmu* dominates tuft cell responses in the distal SI, related to Figure 2**

A) Gating strategy for the identification of tuft cells in the epithelium of the distal SI by flow cytometry (Live CD45^-^ EpCAM^+^ SiglecF^+^). (B) Gating strategy for the identification of ILC2s in the lamina propria (LP) of the distal SI by flow cytometry in SMART13 mice (Live CD45^+^ CD127^+^ KLRG1^+^ IL-13^+^). (C) Gating strategy for the identification of Th1, Th2, and Th17 cells in the LP by flow cytometry: Live CD45^+^ CD3^+^ CD4^+^ cells that are IFNγ ^+^ (Th1), GATA3^+^ (Th2), or IL-17^+^ (Th17). (D) Protist quantification in each region of the intestine by qPCR, in SPF mice singly colonized with each protist. (E) Tuft cell frequency in the epithelium of mice colonized by each protist individually or colonized by both species (Co-col). (F) Protist quantification from *Tmu* and *Tc* co-colonized mice in each region of the intestine. Data points are average of at least 5 mice per group (D, F). Error bars show standard deviation (D, F). *p<0.05, **p<0.01, ***p<0.001, NS, not significant by Mann-Whitney test (D, F) or Student’s t test (E).

**Figure S3: *Tmu* and *Tc* are sufficient to stimulate Th1 and Th17 immunity through an adhesion-independent manner, related to Figure 3**.

(A) Frequency (top) and absolute abundance (bottom) of IFNγ, IL17, and IFNy-IL17 double positive cells in the colonic lamina propria of gnotobiotic mice monocolonized with each protist for three weeks, or left germ free (GF). (B) Abundance of regulatory T cells in the colonic lamina propria of wild type SPF mice colonized with each protist, or uncolonized mice (*Ctrl*). (C) Western Blot of *Tmu* and *Tc* lysates stained with the anti-*Tmu* polyclonal antibody. (D) Representative image of a *Tmu* trophozoite (left) and pseudocyst (right). Protists stained with anti-*Tmu* antibody (cyan), nuclei stained with DAPI (blue). Scale bars are 5µm. (E) Representative image of a *Tmu* cell adhered to the colonic epithelium. Scale bar is 10µm. (F, G) Representative image of a flushed colon from *Tmu* (F) and *Tc* (G) colonized mice. *Tmu* and *Tc* stained with anti-*Tmu* antibody (cyan or magenta, respectively), mucus stained with UEA1 (green), nuclei stained with DAPI (blue), epithelium stained with E-Cadherin (white). Scale bars are 150µm. Each symbol represents an individual mouse (A, B). *p<0.05, **p<0.01, ***p<0.001, ****p<0.0001, NS, not significant by Students t test.

**Figure S4: Characterization of fermentative metabolic pathways in parabasalids, related to Figure 4**.

(A) Tuft cell frequency in the epithelium of the distal small intestine in WT or *Gpr91*^-/-^ mice colonized by each protist or fed succinate in the drinking water. (B) Quantification of extracellular lactate in the cecal contents of mice colonized with each protist for two weeks. (C) Genome assembly statistics for *Tmu* and *Tc* draft genomes. (D) Short chain fatty acid quantification in purified protist lysates (E) Heatmap showing the abundance of 48 metabolites identified in *Tmu* and *Tc* lysates using LC-MS methodologies. Values are plotted as log_2_ of the relative metabolite abundance. (F) Succinate (left) and lactate (right) quantification in culture supernatants of *P. hominis* (order *Trichomonadida*) and *M. colubrorum* (order *Tritrichomonadida*). Data is plotted as mean with SD in B and F. Each symbol represents an individual mouse pooled from two experiments (A). Error bars represent standard deviation (B, F). *p<0.05, **p<0.01, ***p<0.001, NS, not significant by Student’s t test.

**Figure S5. *Tmu* and *Tc* occupy different nutritional niches within the microbiota, related to Figure 5**.

(A) Abundance of *Tmu* and (B) *Tc* in the cecal contents of mice fed diets with different fiber compositions for two weeks. Fibers in the defined diets are limited to inulin (I), cellulose (C), or no fiber (None). (C and D) Relative abundance, from 16S rRNA Sequencing, of the 15 most abundant operational taxonomic units in the distal SI (C) or cecum (D) of mice with or without each protist. Averages of five mice per group are shown. (E) Relative abundance of *A. muciniphila*, as measured by qPCR, in the stool of mice colonized with or without each protist for three weeks. (F) Abundance of genes encoding CAZymes in glycoside hydrolase (GH) families associated with activity on host glycans or plant glycans. White indicates no genes in the indicated GH family, black indicates greater than or equal to 20 genes. Data is plotted as mean in (A, B, and E), with each symbol representing an individual mouse. *p<0.05, **p<0.01, NS, not significant by Mann-Whitney test.

**Figure S6. Fiber deprivation causes *Tmu* to switch to a mucolytic metabolism, related to Figure 6**.

(A) Serial passaging of *Tmu* in PBF medium. Drops in culture density show when *Tmu* was subcultured into fresh media. (B) Survival of *Tc* in PBF medium after 24 hours, with maltose or mucus added as the carbon source. (C) *Tmu* abundance in the stool of mice after two weeks of feeding on def-C chow and different antibiotic treatments. (D) Representative image of *Tmu* in the colon of a mouse fed def-C chow and fed ampicillin in the drinking water. *Tmu* is stained with anti-*Tmu* antibody (cyan), mucus is stained with UEA1 (green), nuclei are stained with DAPI (blue). Scale bar is 100µm. (E) Quantification of *Tmu* localization to the mucus layer in the colons of mice fed complex chow or def-C chow with ampicillin treatment. Higher mucus/lumen ratio indicates tighter localization to the colonic mucus layer. (F) Quantification of total bacterial abundance in the stool of *Tmu*-colonized mice fed def-C chow and given different antibiotic treatments in the drinking water (mice from Figure 6G). Graphs (A, B, C, and D) depict mean. Error bars represent standard deviation (B). Each symbol represents an individual mouse (C, E, F) *p<0.05, **p<0.01, NS, not significant by Mann-Whitney test (C, F) or Students t test (B, E).

**Figure S7. *Tmu* utilizes specific dietary fibers as a carbon source, Related to Figure 7**

(A) *Tmu in vitro* growth curves with different dietary fibers in PBF medium. Dashed lines show maximum growth over 5 days in PBF medium without a supplemented carbon source. (B) Tuft cell frequency and (C) Th2 cell frequency in the distal SI of mice colonized with *Tmu* and fed def-C chow for three weeks. (D) Succinate quantification in the culture supernatants of *Tmu* cultured with maltose or mucus as a carbon source. Graphs show mean with SD (A, D). Each symbol represents an individual mouse (B, C). NS, not significant by Student’s t test.

**Table S1: Reference parabasalid ITS sequences used for identification of parabasalids in human metagenomic datasets, related to Figure 1**.

