## Supplementary figures and images for "Metabolic diversity in commensal protists regulates intestinal immunity and trans-kingdom competition"

### Supplemental Figures

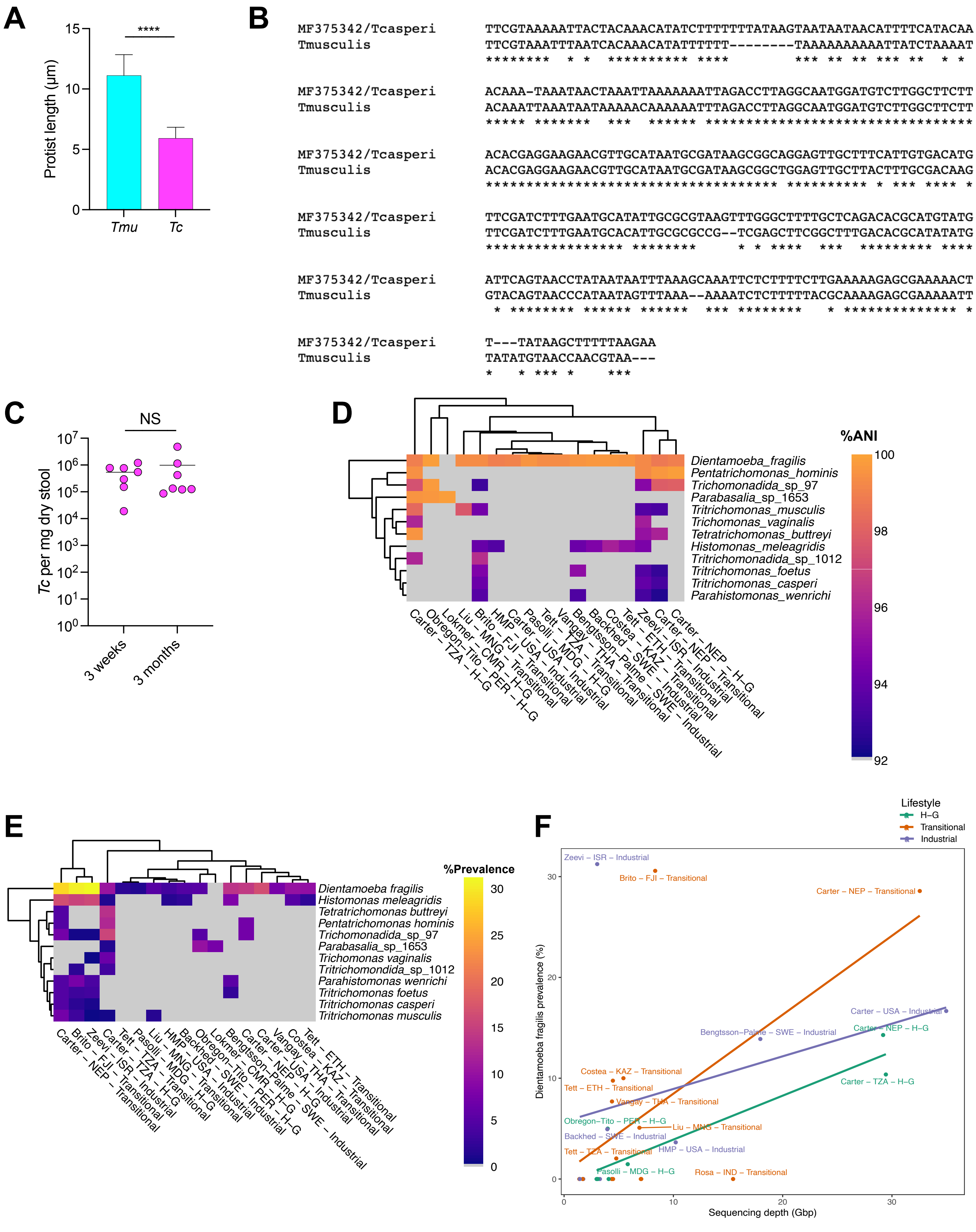

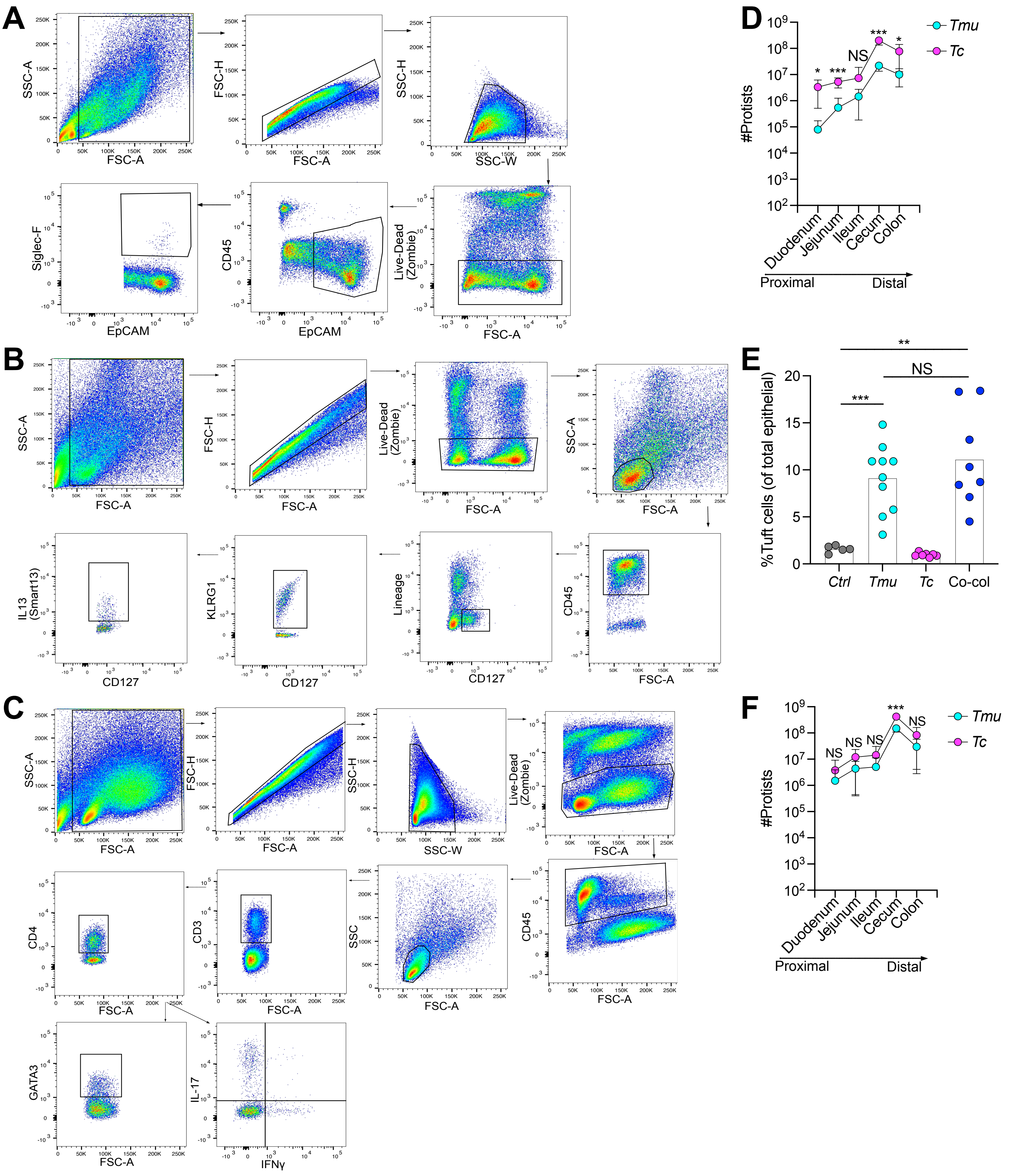

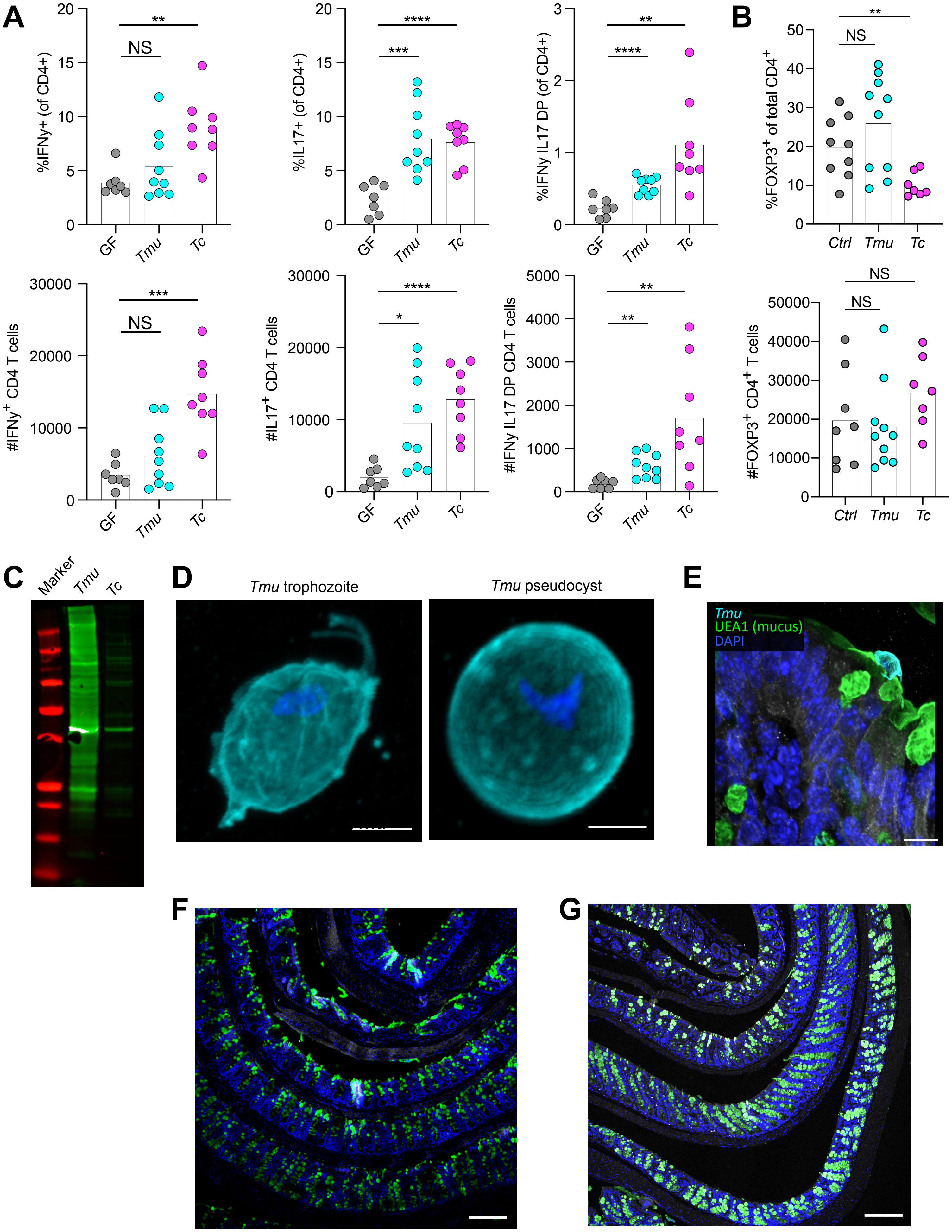

#### Figure S4

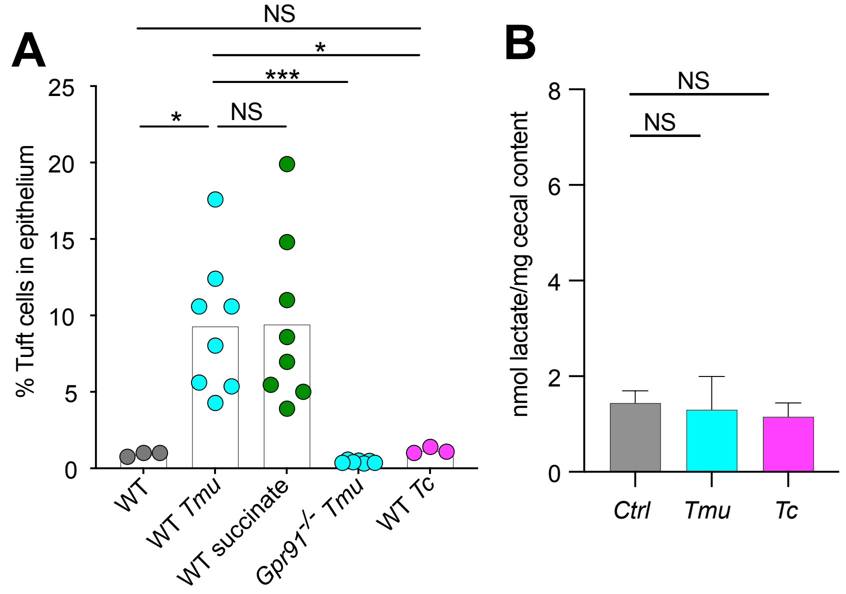


C

| | *T. musculis* | *T. casperi* |
|---|---|---|
| Genome length (Mbp) | 157.8 | 64.3 |
| Gene count | 45,134 | 22,632 |
| $N_{50}$ (Mbp) | 1.477 | 1.213 |
| Coverage | 56.5 | 66.5 |

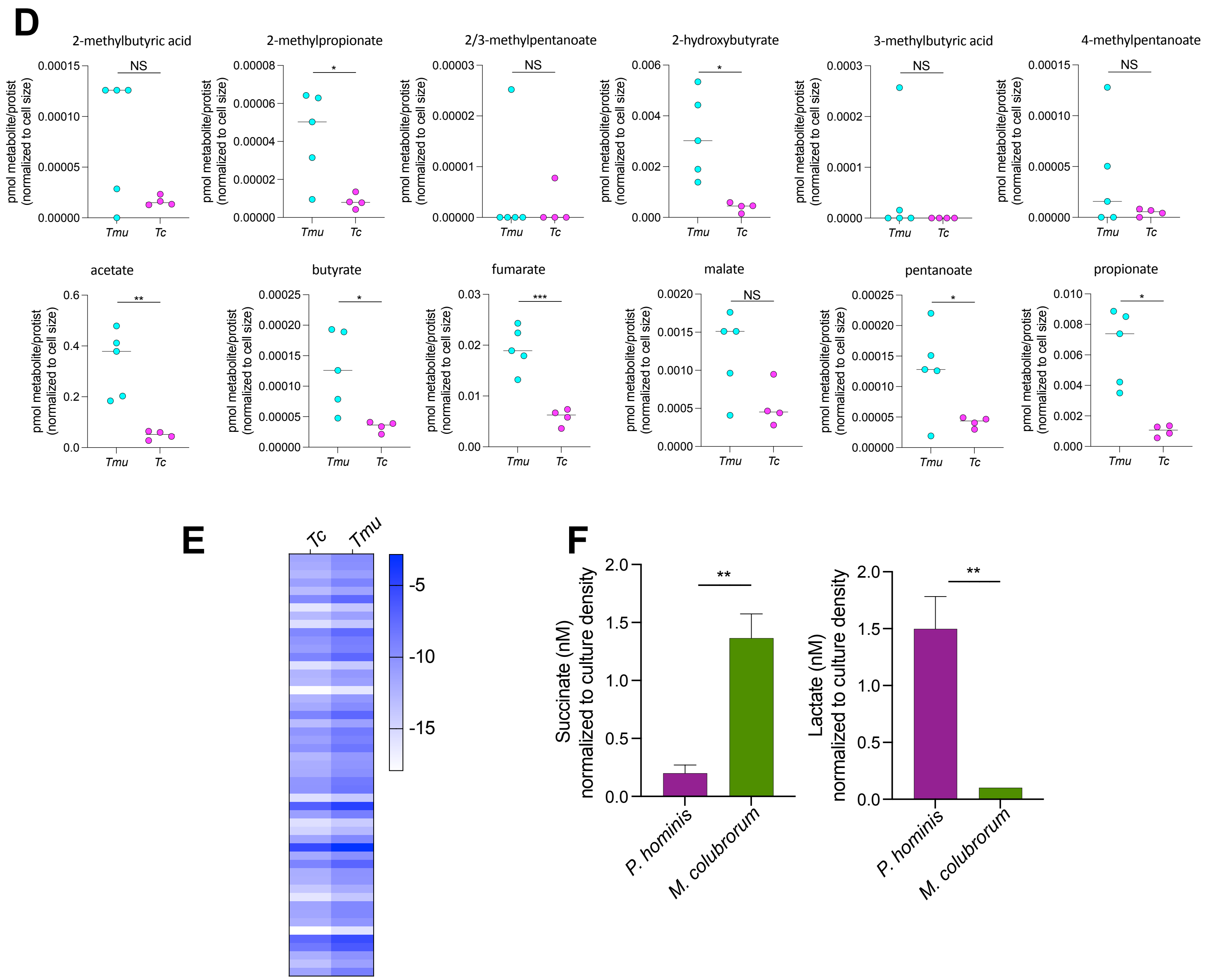

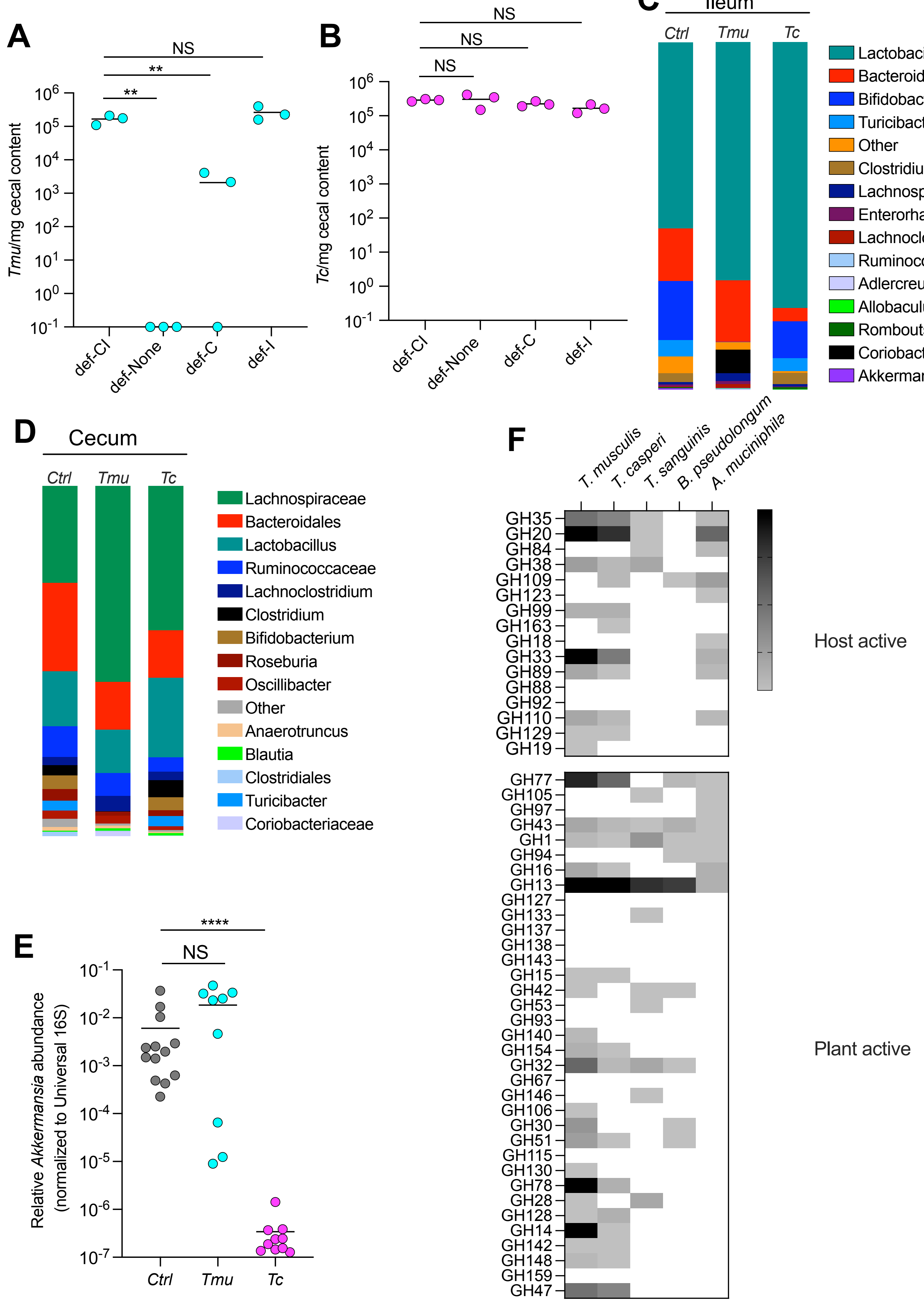

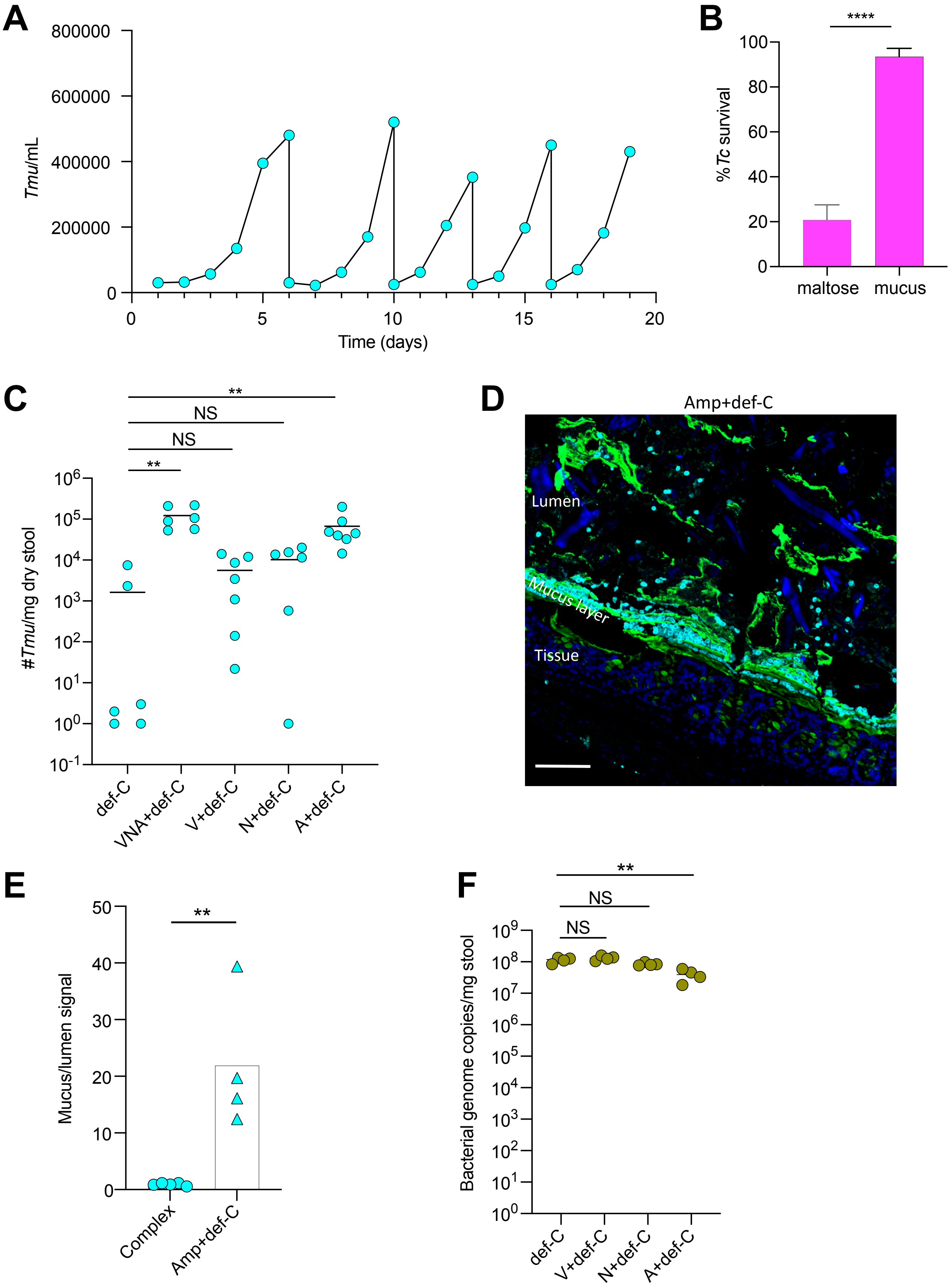

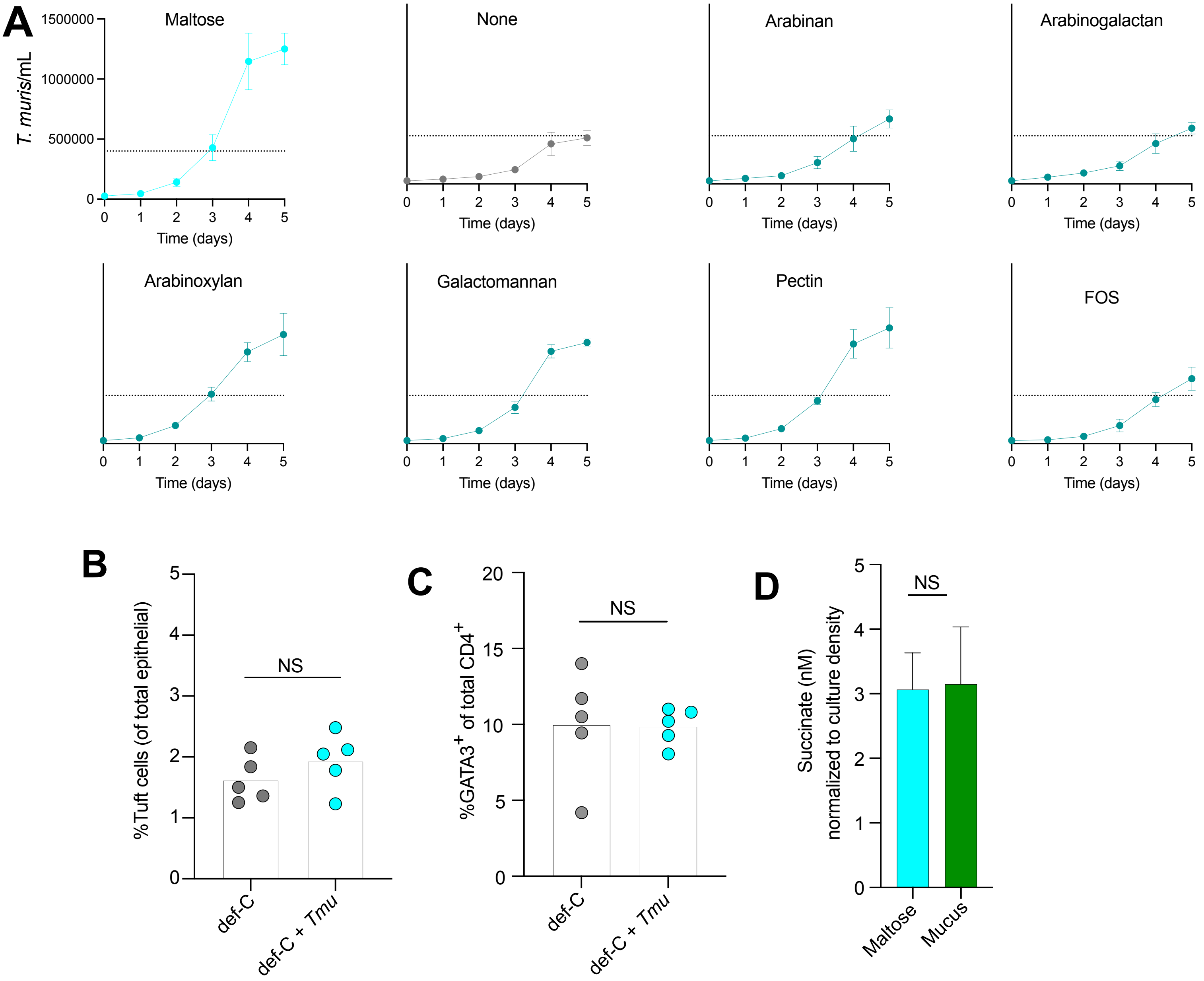
